# Phenotypic melanoma heterogeneity is regulated through cell-matrix interaction-dependent changes in tumor microarchitecture

**DOI:** 10.1101/2020.06.09.141747

**Authors:** Loredana Spoerri, Crystal A. Tonnessen-Murray, Gency Gunasingh, David S. Hill, Kimberley A. Beaumont, Russell J. Jurek, Jagat Chauhan, Gilles C. Vanwalleghem, Mitchell E. Fane, Sheena M. Daignault-Mill, Nicholas Matigian, Glen M. Boyle, Ethan K. Scott, Aaron G. Smith, Samantha J. Stehbens, Helmut Schaider, Brian Gabrielli, Wolfgang Weninger, Colin R. Goding, Nikolas K. Haass

## Abstract

Phenotypic heterogeneity of cancer cells plays a critical role in shaping treatment response. This type of heterogeneity is organized spatially with specific phenotypes, such as sharply demarcated clusters of proliferating and cell cycle-arrested cells, predominating within discrete domains within a tumor. What determines the occurrence of specific tumor cell phenotypes in distinct microdomains of solid cancers is poorly understood. Here, we show that in melanoma spatial organization of phenotypic heterogeneity is dictated by the expression and activity of MITF. We reveal that this lineage survival oncogene controls ECM composition and organization, and ROCK-driven mechanotransduction through focal adhesion maturation and actin cytoskeleton functionality. In turn, altered tumor microarchitecture and structural integrity impact tumor solid stress which then mediates phenotypic heterogeneity through p27^Kip1^. Rho-ROCK-myosin signaling is necessary to transmit the effect of the reciprocal cell-ECM regulation into phenotypic heterogeneity. Our findings place cell-ECM crosstalk as a central driver of phenotypic tumor heterogeneity.

**Significance:** Phenotypic heterogeneity is a major culprit of cancer therapy failure. We demonstrate that phenotypic heterogeneity is controlled through tumor cell-ECM crosstalk resulting in altered tumor microarchitecture, mechanotransduction and Rho-ROCK-myosin signaling. Melanoma shares these physical properties with any solid cancer underscoring the importance of our findings for therapeutically targeting this phenomenon.

## Introduction

Despite the unprecedented success of targeted and immuno-therapies, many patients with advanced cancer still die due to therapy resistance (1,2). Genetic and non-genetic tumor heterogeneity are primary characteristics of solid cancers and are a main cause of promoting drug resistance (3–6). Phenotypic tumor heterogeneity is defined by the coexistence of cells with distinct features within a tumor relating to variations of the epigenome, transcriptome and/or proteome amongst cells (6).

Melanoma is amongst the most genetically and phenotypically heterogeneous cancers at an inter-patient, inter-tumor and intra-tumor level (6). In melanoma, the microphthalmia-associated transcription factor (MITF) has been implicated in tumor heterogeneity and therapy resistance (7–9). MITF is the master regulator of melanocyte biology regulating melanocyte survival, proliferation and melanogenic protein expression and is one of the key factors implicated in the regulation of melanoma cell proliferation and invasion (10–12). MITF is expressed in 56% of human primary melanomas and 23% of melanoma metastases (13). Together with mutant BRAF, amplified MITF can act as an oncogene (14). Additionally, a recurrent mutation in MITF (E318K) predisposes to familial and sporadic melanoma (15,16). MITF both enhances and inhibits melanoma proliferation by controlling cell cycle regulators. These conflicting functions can be explained by a rheostat model, which states that *very high* levels and activity of MITF predispose melanoma cells to cell cycle arrest and differentiation, whereas *critically low* MITF levels lead to cell cycle arrest, senescence and apoptosis; only intermediate levels favor proliferation (17–21). Two distinct gene expression signatures in melanoma are characterized by MITF levels: The *intermediate-high* MITF (from here referred to as MITF^high^) proliferative phenotype and the *intermediate-low* MITF (from here referred to as MITF^low^) invasive phenotype (22), which oscillate *in vivo* and underlie the phenotype switching phenomenon (23). These findings demonstrate that MITF is closely tied with melanoma progression, but its precise role remains unclear. MITF expression has been associated with therapy response, with both beneficial and detrimental effects identified (24). The MITF rheostat model is likely to be central to divergent responses to MAPK pathway inhibition, with intermediate MITF expression serving as a marker for treatment response while very low or very high MITF levels conferring resistance. In this context, it is plausible to attribute the underlying efficacy of drug response to MITF-mediated regulation of the cell cycle. In fact, G_1_-arrested and dormant cells are more prone than cycling cells to develop drug tolerance and resistance (25–27), while excessive proliferation may counteract drug-mediated cell killing, ultimately hindering drug efficacy. Intermediate levels of MITF activity may provide an effective therapeutic window with a balance in proliferation and G_1_-arrest in between the two extreme scenarios (3).

In addition to genetic and epigenetic factors, the microenvironment plays a pivotal role in determining the cycling state of a cell. Beyond well-known determinants of proliferation and quiescence such as nutrients, growth factors and mitogens, the extracellular matrix (ECM) is a key contributing factor to regulating cell proliferation, both in normal tissue and in cancer (28,29). With its multitude of biochemical and biomechanical properties, the ECM not only acts as a scaffold to determine tissue morphology and physical characteristics, but also influences signaling and biology of the cells that it surrounds (30,31). Cell-matrix adhesions are crucial mediators of biochemical and mechanical cues between the ECM and cells, as they transmit information of the composition as well as the rigidity of the ECM (32). This subsequently triggers a multitude of signaling cascades that result in adaptive responses affecting proliferation, morphology, motility, matrix deposition and organization and many other cellular properties, including therapy resistance (33).

Here we demonstrate that phenotypic melanoma cell heterogeneity is decreased *in vitro* and *in vivo* by increased MITF expression and activity. This phenomenon is not associated with cell cycle profile changes in adherent culture, suggesting that MITF mediates these effects through factors requiring a 3D environment, rather than simply through its rheostat-based influence on proliferation. We show that MITF expression affects reciprocal cell-ECM interactions which in turn, through ROCK signaling impact solid stress, p27^Kip1^ regulation and ultimately intratumor phenotypic heterogeneity. This microenvironment-heterogenity control axis may act in other solid cancers where the ECM has been shown to influence cancer progression and response to therapy (34,35).

## Results

### Phenotypic heterogeneity in melanoma is dependent on MITF level and activity

We investigated the role of MITF in phenotypic heterogeneity using the intratumoral distribution of differentially cycling melanoma cells *in vivo* as a read out. Melanoma cell lines were transduced with the fluorescent ubiquitination-based cell cycle indicator (FUCCI) system (36,37) to differentially visualize the G1 (monomeric Kusabira Orange2, mKO2; displayed in magenta) and S/G2/M (monomeric Azami Green, mAG; green) phases of the cell cycle. Endogenously MITF^low^ (1205Lu and C8161) and MITF^high^ (WM983C, WM164, 451Lu) FUCCI-transduced melanoma cell lines (Figure 1A) were xenografted subcutaneously into NOD/SCID mice. MITF and its transcriptional target RAB27A (38) exhibited a similar expression pattern, indicating that MITF was functional in the MITF^high^ cell lines (Figure 1A). Analysis of histologic sections of these tumors by confocal microscopy revealed two distinct cohorts based on distribution of differentially cycling cells (Figure 1B, Figure S 1A). The MITF^high^ cohort exhibited a predominantly homogeneous distribution of mKO2- and mAG-positive cells, indicating proliferation throughout. In contrast, the MITF^low^ cohort showed sharply demarcated proliferating and G_1_-arrested clusters, predominantly composed of mKO2^+^ cells (Figure 1B, Figure S 1A).

**Figure 1:**
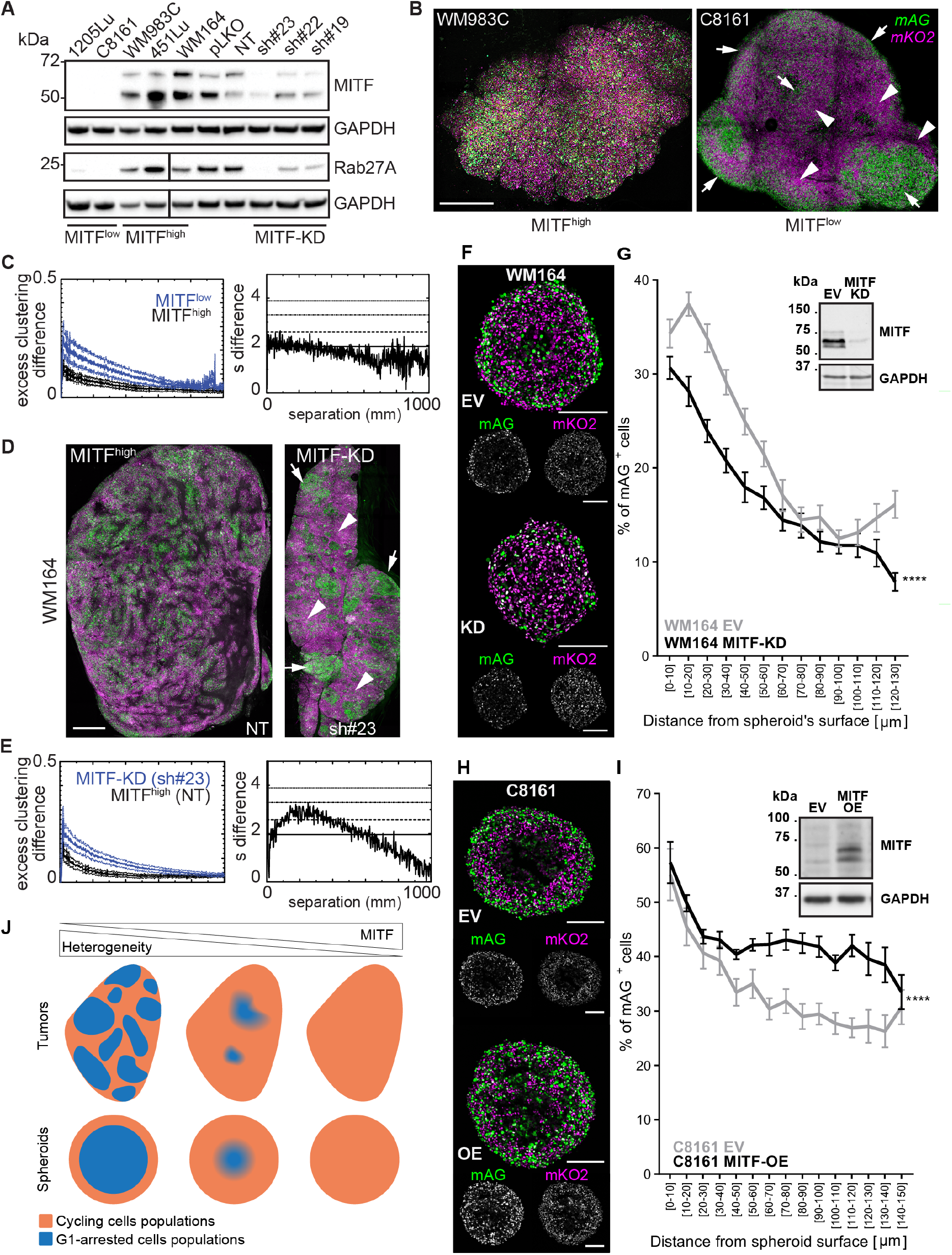
Phenotypic heterogeneity in melanoma is dependent on MITF level and activity. **(A)** Protein levels of MITF and its downstream effector RAB27A in the melanoma cell lines used for engrafting. Immunoblots are representative of three independent experiments. The vertical line indicates an unrelated excised lane from the original blot. **(B,D)** Representative examples of predominantly homogeneously proliferating MITF^high^ (WM983C (B) and WM164-NT (D)) and clustered MITF^low^ (C8161 (B) and WM164-sh#23 (D)) xenografts. Arrows and arrowheads indicate proliferating and G_1_-arrested clusters, respectively; scale bars: 1 mm. **(C,E)** Quantitative analysis of the differences in mKO2^+^ (magenta) and mAG^+^ (green) cell clustering in the two cohorts. Clustering of mKO2^+^ and mAG^+^ cells was first compared to a random distribution, then the absolute clustering differences were obtained for each tumor image. Error bars represent ±SEM of at least 19 (C) or 14 (E) tumors per cohort. In the sigma difference (s difference) plots, solid, dashed, dot-dash and dotted horizontal lines correspond to p-values of 0.05, 0.01, 0.001, 0.0001. **(F,H)** Representative FUCCI fluorescence images (monomeric Azami green, mAG, green; monomeric Kusabira Orange2, mKO2, magenta) of equatorial cryosections of spheroids with MITF depleted (WM164^MITF-KD^) or overexpressed (C8161^MITF-OE^), and the respective controls (WM164^EV^ and C8161^EV^). Areas of mAG^+^ and mKO2^+^ cells proliferate, while zones depleted of mAG^+^ cells are G_1_-arrested; scale bars: 200 μm. **(G,I)** Analysis of the spatial distribution of the proliferation area. Graphs show the percentage of mAG^+^ cells (mAG^+^/(mAG^+^ + mKO2^+^)) as a function of distance from the spheroid surface. Note, cells beyond specific spheroid depths (cell-line dependent) were omitted from the analysis because of high variability driven by core necrosis, characterized by absence of FUCCI fluorescence signal. WM164: n=16 (EV), n=16 (MITF-KD); C8161: n=11 (EV), n=12 (MITF-OE); data: mean ±SEM; analysis: two-way ANOVA for the effect of MITF on % of mAG^+^ cells; **** p<0.0001. Immunoblots show the level of MITF knockdown (B) and overexpression (D). **(J)** Schematic of how intratumoural heterogeneity manifests in tumours and in spheroids. NT, non-targeting shRNA, mKO2, monomeric Kusabira Orange2, magenta; mAG, monomeric Azami Green, green; KD, knock-down; EV, empty vector; OE, overexpression.

To objectively analyze the distribution of proliferating cells within these xenografts, we adapted an astrophysical correlation function that has been used to describe the clustering of galaxies (39). We measured the number of mKO2^+^ or mAG^+^ cell pairs at a given separation and compared that to the number of pairs we would expect if the cells were randomly distributed. To identify clustering differences irrespective of which cells (mKO2^+^ or mAG^+^) were more strongly clustered, we plotted the absolute difference in the mKO2^+^ and mAG^+^ cell clustering (Figure 1C, Figure S 1B-E). This analysis confirmed that the MITF^low^ cohort had significantly greater clustering compared to the MITF^high^ cohort (Figure 1C).

To assess the functional role of MITF in intratumor phenotypic heterogeneity, we depleted MITF in endogenously MITF^high^ FUCCI-WM164 melanoma cells using shRNA-expressing lentiviral particles (MITF-KD) and generated xenografts. Downregulation of MITF protein levels by approximately 80% (Figure 1A; Figure S 1F) did not affect the cell cycle profile in 2D culture (Figure S 1G).

While xenografts expressing non-targeting shRNA (NT) controls showed a more homogeneous cell cycle pattern, MITF-KD xenografts displayed proliferating and G_1_-arrested cell clusters (Figure 1D, Figure S 1H). Quantitative clustering analysis confirmed a significant difference between mKO2^+^ and mAG^+^ cell clustering in MITF^high/shRNA^ xenografts compared to MITF^high/NT^ controls (Figure 1E). We thus conclude that the presence of G_1_-arrested clusters in melanoma xenografts, indicative of phenotypic heterogeneity, is linked to low MITF levels.

To elucidate the mechanism underlying MITF-mediated phenotypic heterogeneity, we next utilized a 3D FUCCI melanoma spheroid model (37,40). Melanoma spheroids recapitulate the tumor microenvironment and cell cycle behavior in relation to oxygen and nutrient availability observed in xenografts (37,40), except that these are supplied by diffusion and not through vasculature, resulting in a uniform and concentric oxygen and nutrient gradient (41). This leads to proliferation in the spheroid periphery, G_1_-arrest in the center and, in some cases, a necrotic core (37); the width of these zones is cell line-specific.

As melanoma cells have a high mutation rate, we proceeded to use isogenic cell lines to assess the mechanistic relationship of MITF expression and cell cycle distribution. In addition to WM164, we depleted MITF in an additional MITF^high^ melanoma cell line (WM983B) (Figure 1F,G; Figure S 2A,B) and over-expressed MITF in two endogenously MITF^low^ melanoma cell lines (C8161 and WM793B) (Figure 1H,I; Figure S 2C,D). We then analyzed the spatial distribution of mKO2^+^ and mAG^+^ cells in spheroids using confocal microscopy. Because of the relatively uniform and concentric gradient of cell cycle activity, we measured % mAG^+^ (mAG^+^/(mAG^+^ + mKO2^+^)) cells in relation to the distance from the spheroid surface (42). Both empty vector control (EV) and MITF-KD spheroids displayed an external ring of proliferating cells and a G_1_-arrested center. However, the G1-arrested center significantly extended to almost reach the surface of the MITF-KD spheroids (Figure 1F; Figure S 2A). We confirmed this difference by calculating the percentage of mAG^+^ cells, which was lower in MITF-KD spheroids when plotted as a function of distance from the surface (Figure 1G; Figure S 2B). Analogously, ectopically over-expressing MITF in endogenously MITF^low^ cell lines (MITF-OE) reduced the G1-arrested spheroid center resulting in a wider proliferating periphery compared to controls (Figure 1H,I; Figure S 2C,D). These data support an inverse relation between MITF expression and the extent of G_1_-arrested zones *in vivo* and *in vitro*. Of note, similar to MITF depletion, MITF overexpression had little or no effect on the cell cycle profile in 2D culture (Figure S 1H), confirming that our observation is a 3D-specific phenomenon. We define phenotypic heterogeneity in xenografts as sharply demarcated proliferating and G1-arrested clusters, which is reflected in the spheroids by sharp demarcation of the G1-arrested center (Figure 1J). The definition of “homogenous” would be 100% cycling cells throughout, which we never observe in reality. Thus, tumors and spheroids are more homogenous when MITF is high compared to low, where they are more heterogeneous (Figure 1J). As both MITF levels and tumor cell heterogeneity are important for melanoma therapy outcome (3), we next investigated the molecular basis of MITF-controlled phenotypic heterogeneity in an effort to better understand and potentially control it.

### MITF overexpression decreases structural integrity and solid stress in melanoma spheroids

Alteration of MITF levels not only affected phenotypic heterogeneity but also spheroid morphology. MITF-OE melanoma spheroids appeared consistently larger than control spheroids (EV), while MITF-KD spheroids appeared reduced in size (Figure 1F,H; Figure S 2). These observations were confirmed by bright field microscopy measurement of spheroid projected areas (Figure S 3A). We then imaged spheroids using single plane illumination microscopy (SPIM) to measure the equatorial and polar diameters (Figure 2A,B; Figure S 3B,C) and to assess overall spheroid shape. We revealed a consistently decreased roundness (equatorial to polar diameter ratio) in spheroids from three different cell lines upon MITF overexpression (Figure 2C). In contrast, MITF-KD had no significant effect on roundness (Figure 2C), likely due to the fact that a spheroid that is already round (roundness in average between 0.55 and 0.80 for endogenously MITF^high^ spheroids (Figure 2C), is physically limited to change towards a rounder morphology.

**Figure 2:**
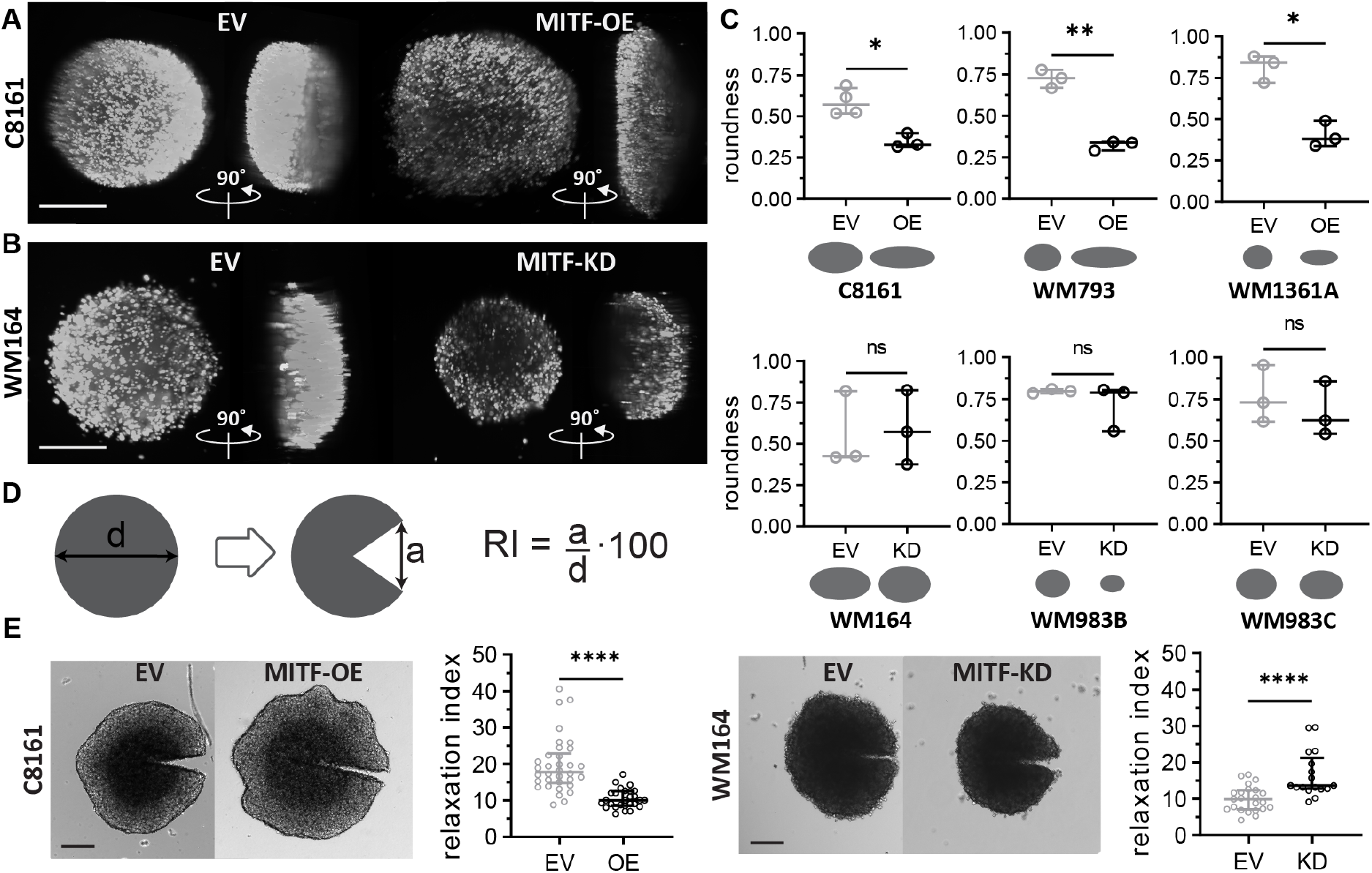
MITF overexpression decreases structural integrity and solid stress in melanoma spheroids. Representative SPIM images (utilizing mAG for fluorescence) and quantitation of polar and equatorial diameters of live **(A)** C8161^EV/MITF-OE^ and **(B)** WM164^EV/MITF-KD^ spheroids. Magenta and cyan lines indicate measured equatorial and polar diameters, respectively, scale bars: 200 μm. **(C)** Spheroid roundness calculated as the ratio of equatorial to polar diameter and schematics showing spheroid silhouettes from a side-on view, demonstrating changes in spheroid size and roundness. **(D)** Schematic of solid stress assessment protocol and formula to calculate the relaxation index (RI). **(E)** Representative transmitted-light microscopy images of incised C8161^EV/MITF-OE^ and WM164EV^/MITF-KD^ spheroids and quantitation of their respective relaxation index. Scatter dot plot of median and interquartile range; analysis: paired t-test (C) and Mann Whitney test (E), ****p<0.0001, ***p<0.001, ** p<0.01, * p<0.05, ns p>0.05.

To further assess the physical properties, we asked whether the morphological changes were associated with altered solid stress, i.e. the mechanical stress that is contained in and transmitted by the solid and elastic elements of the ECM and cells in the tumor microenvironment (43). To address this question, we incised spheroids to approximately 50% of their diameter and measured the incision opening to calculate the relaxation index (RI) indicative of solid stress (Figure 2D) (44–46). MITF overexpression significantly decreased the relaxation index of C8161 spheroids, supporting a reduced solid stress (Figure 2E). Analogously, MITF knock-down of WM164 spheroids increased solid stress (Figure 2E), despite the fact that it did not significantly alter their morphology (Figure 2B). We demonstrate that high MITF levels result in flattening and solid stress reduction of melanoma spheroids, while cell density is not affected. These findings strongly suggest that MITF regulates factors involved in altering both architecture and solid stress in melanoma spheroids.

### MITF expression affects molecular processes involved in ECM organization

To decipher the molecular mechanisms underlying, and potentially linking, the MITF-driven changes in architecture, solid stress and phenotypic heterogeneity, we performed a proteomics study based on MITF expression (C8161^MITF-OE^ compared to C8161^EV^ spheroids and WM164^EV^ compared to WM164^MITF-KD^ spheroids) to identify differentially expressed proteins (Figure 3A). A total of 66 and 33 proteins were found to have statistically significant different levels between EV/OE and EV/KD, respectively. The STRING database was used to build a protein-protein network from the differentially expressed proteins and gene ontology enrichment was assessed on this network. Amongst the identified processes, ECM organization emerged as the predominant hit with a highly significant FDR value and presence in both cohorts of comparison (Figure 3B). Next, we sought to validate the hits that enriched these processes (Figure 3C) as well as other proteins that showed significant changes in our proteomics study by immunoblotting (Figure 3D).

**Figure 3:**
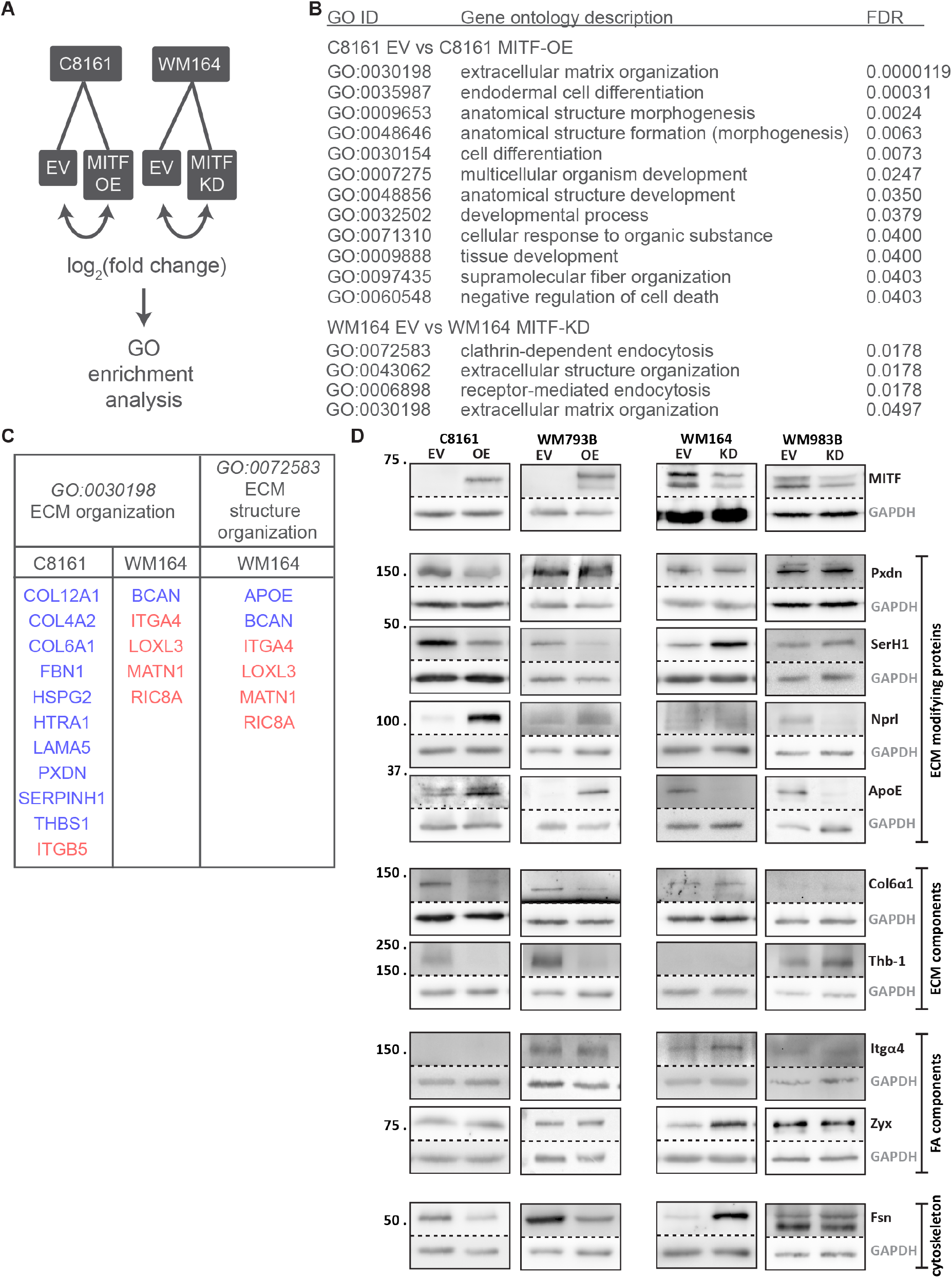
MITF expression affects levels of proteins involved in ECM and anatomical structure. **(A)** Schematic of proteomics comparisons and downstream analysis. Lysis of whole (non-dissociated) spheroids assured that samples contained both proteins found within cells as well as within the ECM secreted by the cells. **(B)** Gene Ontology pathways identified through STRING-based pathway enrichment analysis. **(C)** List of the hits that were enriched the ECM organization pathways. Blue: increase in MITF-OE/KD compared to EV; red: decrease in MITF-OE/KD compared to EV. **(D)** ECM interaction protein immunoblots of spheroid lysates.

In C8161^MITF-OE^ compared to C8161^EV^ spheroids, we confirmed a decrease in peroxidasin, an enzyme responsible for fibrillar network assembly of collagen IV, fibronectin and laminin (47,48), and a decrease in serpin H1, another crucial enzyme for collagen maturation (49,50). We found higher levels of neprilysin, an enzyme that degrades elastin (51), and of apolipoprotein E, an ECM-secreted protein that suppresses the expression of extracellular matrix genes such as collagen I and lysyl oxidase, and whose loss correlates with focal adhesion kinase (FAK) activation in cell-matrix adhesion (52,53). Furthermore, the level of thrombospondin-1, an ECM-secreted protein that increases collagen expression, matrix organization and focal adhesion maturation (54–56), was decreased upon MITF overexpression. Reduced collagen VI-α1 levels was in line with these observations. There was also a decrease in level of fascin, an actin-bundling protein whose activity is regulated by cell-matrix adhesion, including attachment to thrombospondin-1 (57,58). Similarly, serpin H1, neprilysin, apolipoprotein E, collagen VI-α1, thrombospondin-1 and fascin changed accordingly in another cell line, WM793, when MITF had been overexpressed (Figure 3D). Altogether, these findings suggest high MITF levels drive a reduction in cell-matrix interactions, in addition to alterations in ECM components and organization.

Considering the opposing effect of MITF overexpression and knockdown on spheroid morphology and solid stress, reverse protein changes were expected upon MITF knockdown. Serpin H1 was indeed increased, and apolipoprotein E decreased in WM164^MITF-KD^ cells compared to WM164^EV^ cells, but peroxidasin, neprilysin and collagen VI-α1 were not affected and collagen IV-α2 decreased (Figure 3D). Levels of two key components of focal adhesions, integrin-α4 and zyxin, were raised upon MITF silencing, in agreement with increased thrombospondin and reduced apolipoprotein E levels. Fascin levels were increased, supporting an increased cytoskeleton functionality. Some of these protein level changes were reflected upon MITF silencing in a different cell line, WM983B (Figure 3D). In summary, the four examined cell lines showed different protein changes amongst them, which nevertheless entailed congruent downstream effects. Protein changes observed upon MITF overexpression indicated decreased focal adhesion maturation and reduced organization of the ECM and the cell’s actin cytoskeleton, whereas MITF silencing resulted in enhancement of these three processes.

To support the occurrence of this MITF-driven regulation of the cell-ECM bidirectional crosstalk at a broader scale, we interrogated mRNA expression datasets from three independent panels of melanoma cell lines, namely the CCLE dataset (Figure 4A), an in-house collection of 12 melanoma cell lines (Figure 4B) and a panel of 53 cell lines clustered according to phenotype (Figure 4C). The pattern of expression observed in the cell lines was broadly reproduced in the clinical samples from the TCGA melanoma cohort ranked by MITF expression (Figure 4D). These datasets supported an inverse correlation between MITF, and the hits identified in our four cell lines that promote cell-ECM crosstalk as well as a positive correlation with apolipoprotein E and neprilysin, two molecular components known to decrease ECM organization.

**Figure 4:**
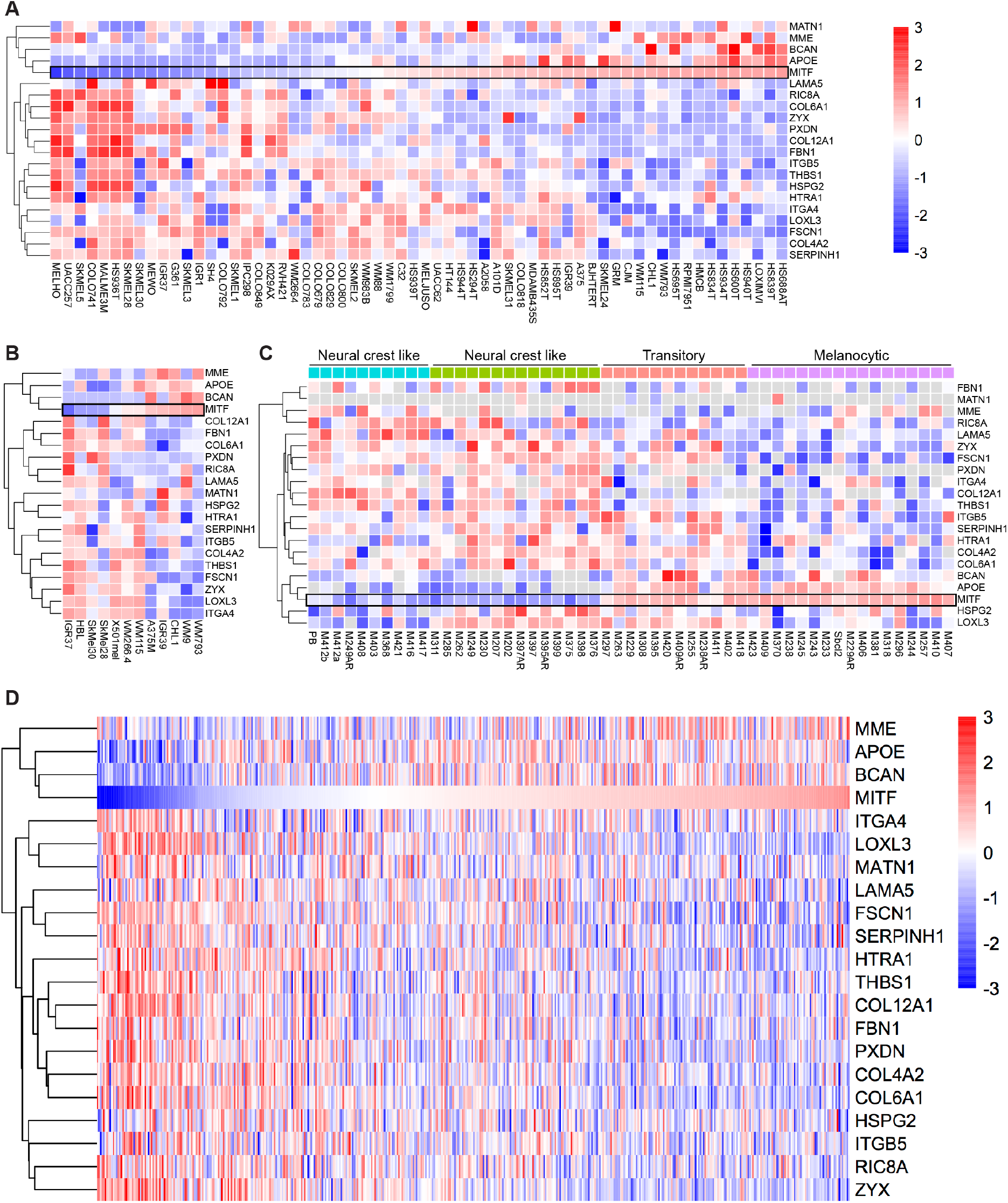
RNA expression heat maps stratified for MITF expression levels. Relative expression of the indicated genes in the (A) Cancer Cell Line Encyclopedia (CCLE) melanoma cell line database, (B) set of melanoma cell lines described by Goding et al. (115), (C) set of melanoma cell lines described by Tsoi et al. (114) and (D) TCGA human melanoma cohort. In all datasets ranking is by expression of MITF (indicated by a black rectangle outline) except for the dataset shown in (C) where the cell lines are grouped according to their phenotype.

Based on ChIP-seq analysis (Figure S 4), MITF binds to *PXDN, MME, HSPG2, LAMA5, MATN1, COL4A2, BCAN, HTRA1, ITGB5, ZYX* and *FSCN1*, indicating that these genes may be direct transcriptional targets of MITF. Lack of binding of MITF to *ITGA4, COL6A1, COL12A1, SERPINH1, APOE, FBN1, THBS1, LOXL3* and *RIC8A* suggests indirect regulation.

### MITF expression levels affect central components of ROCK-driven cell-ECM interaction

To answer whether MITF expression levels affect ECM-mediated signaling, we first investigated focal adhesions, which are key multiprotein complexes in this process. We visualized and compared focal adhesion morphology in the two cohorts by immunostaining the cells with an antibody against tyrosine 397 phosphorylated focal adhesion kinase (pFAK) (Figure 5A), a tension-sensitive focal adhesion component that requires actomyosin contractility (59). The number of focal adhesions per cell remained unchanged both upon MITF overexpression and depletion (Figure S 5A,C). However, we noticed alterations in adhesion morphology, resulting in more rounded adhesions in C8161^MITF-OE^ than in C8161^EV^ and more elongated adhesions in WM164^MITF-KD^ compared to WM164^EV^. This was confirmed by quantitation of focal adhesion morphology (Figure 5A, Figure S 5B,D). As actomyosin-induced contractility stabilizes adhesion formation and maturation, less elongated focal adhesions imply a decreased adhesion maturation and reduced actin fiber attachment to focal adhesions, which subsequently weakens cell contractility.

**Figure 5:**
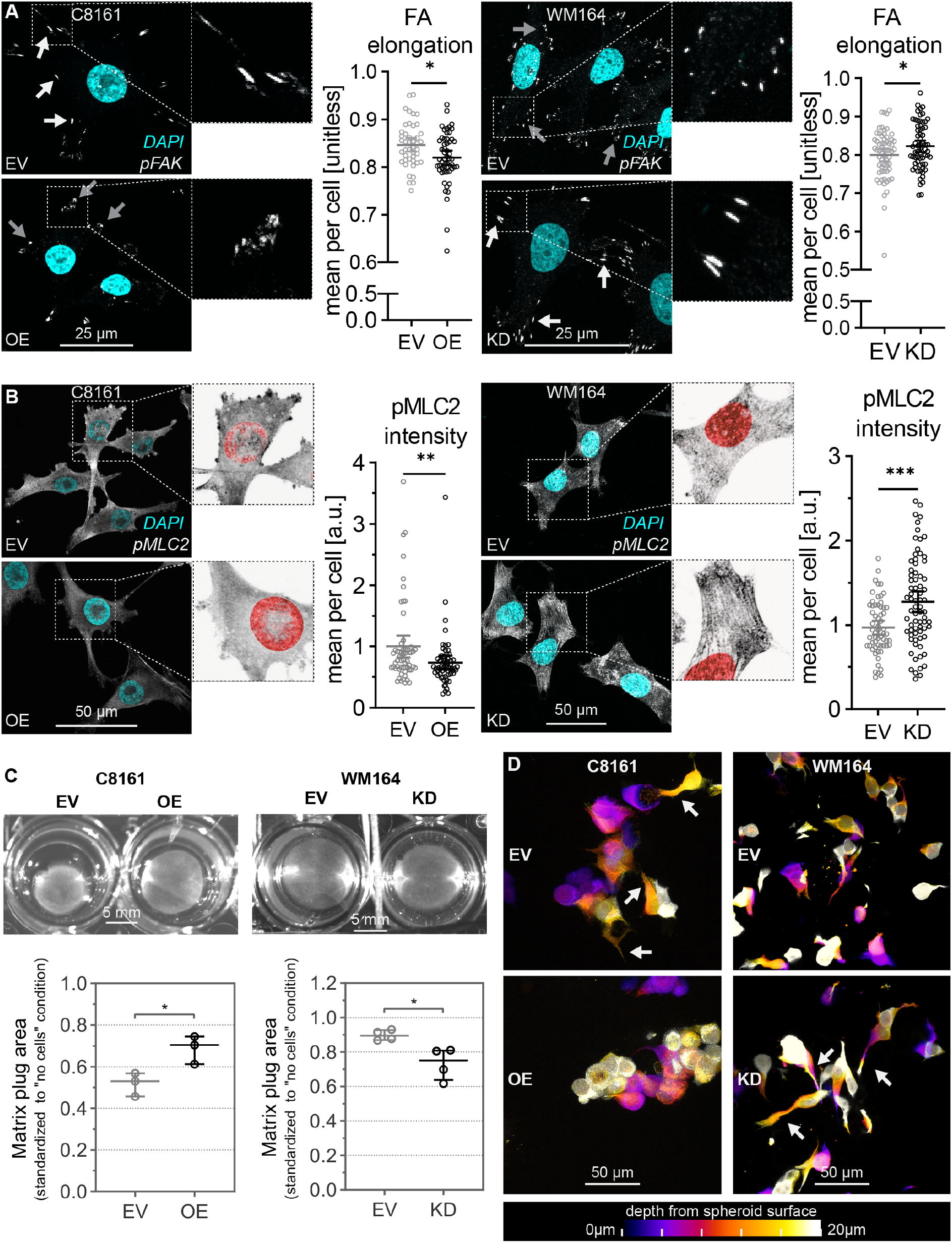
MITF expression levels affect central components of ROCK-driven mechanotransduction. **(A,B)** Representative immunofluorescence microscopy images (maximum intensity projection) of pFAK-Y397 **(A)** and pMLC2-S19 **(B)** detection in C8161^EV^ versus C8161^MITF-OE^ and WM164^EV^ versus WM164^MITF-KD^ cells and quantitation of the mean focal adhesion elongation per cell based on pFAK-Y397 signal **(A)** and pMLC2-S19 mean fluorescence intensity signal per cell and **(B)**. White and gray arrows indicate elongated and rounded focal adhesions, respectively **(A)**. Scatter dot plot of data mean with 95% CI; each dot represents a cell; n>43 cells from three independent experiments; analysis: unpaired t-test (A) and Mann Whitney test (B), * p<0.05, ***p<0.001. FA, focal adhesion. Higher magnification images of pMLC2-stained cells are contrast-inverted to better visualize MLC2 fibers **(B)**. **(C)** Representative images of matrix contraction assay and quantitation of collagen gel plug contraction for C8161 and WM164 cells. Data normalized to an acellular collagen control; scatter dot plot of data median and interquartile range; each dot represents the mean of an independent experiment; n=3 experiments; analysis: paired t-test, * p<0.05. **(D)** Representative Z-stack maximum intensity projection depth color-coded confocal images of C8161^EV^ and C8161^MITF-OE^ and WM164^EV^ and WM164^MITF-KD^ spheroid cells (n=3 independent experiments). Arrows indicate dendrites.

ROCK-driven actomyosin contractility increases tissue stiffness as well as tumor growth and progression (60). Phosphorylation of MLC plays a key role in the Rho-ROCK-myosin pathway, which regulates actin-based machineries that drive mechanosensitive alterations in cell and tissue morphology. This mechanism is known to influence spheroid architecture and compaction (61–63). As a measure for ROCK-driven actomyosin contractility, we assessed the level and sub-cellular pattern of serine 19 phosphorylated myosin light chain 2 (pMLC2) via immunofluorescence in C8161^EV^, C8161^MITF-OE^, WM164^EV^ and WM164^MITF-KD^ cells (Figure 5B). In C8161^MITF-OE^ cells the mean fluorescence signal intensity for pMLC2 was significantly decreased to approximately 75% of C8161^EV^ levels, whereas it was approximately 30% higher in WM164^MITF-KD^ than in WM164^EV^ cells, suggesting an inverse relation between MITF level and contractility in these cells. Furthermore, increased fiber-like pattern of pMLC2 was noticeable in WM164^MITF-KD^ versus WM164^EV^ cells, characteristic of pMLC2 association with actin stress fibers and of contractile structures.

To assess the effect of changed pMLC2 protein levels on the cells’ ability to organize the surrounding matrix, we performed a matrix contraction assay where we embedded C8161^EV^, C8161^MITF-OE^, WM164^EV^ and WM164^MITF-KD^ cells in collagen and measured the matrix plug area as a function of time. Over a 4-day period, C8161^MITF-OE^ cells showed a 35% decrease in their ability to contract the collagen plug compared to C8161^EV^ cells, whereas the WM164^MITF-KD^ cells contracted the collagen plug 62% more effectively than WM164^EV^ (Figure 5C). This indicated that MITF^high^ melanoma cells have an impaired ability to contract the surrounding matrix.

We next studied cell morphology within spheroids. Membrane protrusions play a crucial role in cell-matrix adhesion via bi-directional signaling and physical connections between the ECM and the cell cytoskeleton (64), which in turn are necessary for mechanosensing (65). We generated C8161^EV^, C8161^MITF-OE^, WM164^EV^ and WM164^MITF-KD^ spheroids with unstained cells and cells stained with CellTracker™ in a ratio of 50:1, to allow clearer visualization of cell morphology within 3D spheroids. Images display maximum intensity projection of color-coded depth Z-stacks. Confocal microscopy of CUBIC-cleared spheroids revealed a high degree of dendricity of C8161^EV^ cells when compared to C8161^MITF-OE^ cells and of WM164^MITF-KD^ cells when compared to WM164^EV^ cells (Figure 5D), suggesting an inverse association of MITF level with dendrite projections and ECM adhesion in these cells. Altogether, these findings support the hypothesis of an inverse regulation of ROCK-driven mechanotransduction by MITF.

### MITF affects p27^Kip1^ expression

To further characterize how MITF mediates phenotypic heterogeneity, we asked how changes in spheroid architecture could affect cell cycling and modulate the extent and distribution of G1-arrest in spheroids. Cyclin-dependent kinase inhibitor 1B (p27^Kip1^) prevents the activation of cyclin E-CDK2 and cyclin D-CDK4 complexes, controlling cell cycle progression at G1. Compressive stress in tumor spheroids leads to increased p27^Kip1^ expression resulting in reduced cell proliferation (66). We therefore interrogated the p27^Kip1^ protein level in MITF-OE and MITF-KD cells compared to their respective EV control cells.

The p27^Kip1^ level in C8161^EV^ spheroid progressively increased from the periphery towards the center whereas it remained low, or even decreased, upon MITF overexpression (Figure 6A,B). We attributed the p27^Kip1^ level decline when further approaching the spheroid center in C8161^EV^ spheroids to necrosis. In WM164 spheroids the p27^Kip1^ level progressively increased from the periphery towards the center in both EV and MITF-KD conditions, however it was consistently higher in MITF-KD spheroids (Figure 6C,D).

**Figure 6:**
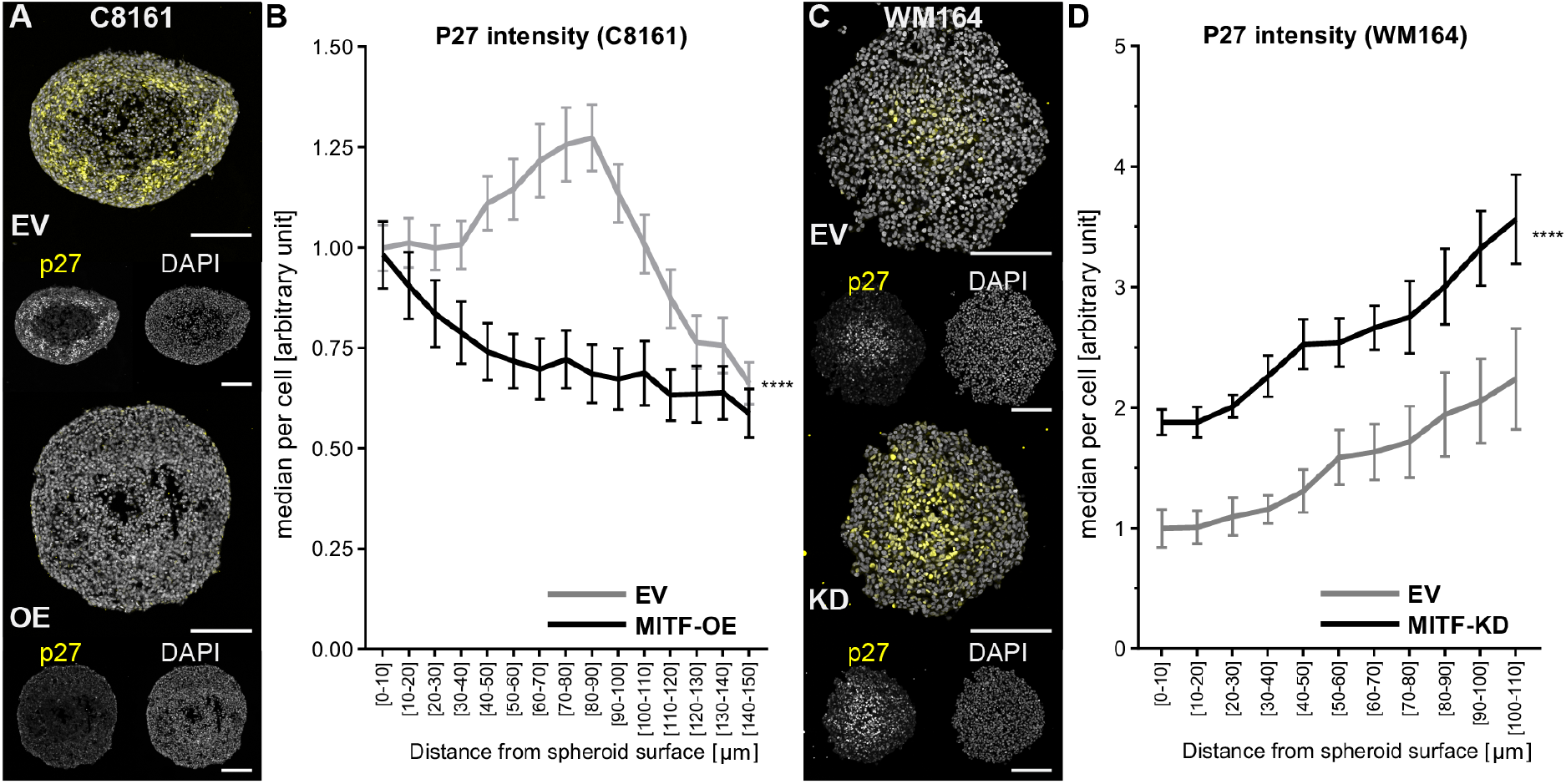
**(A,C)** Representative p27^Kip1^ fluorescence images of equatorial cryosections of spheroids with MITF overexpressed (C8161^MITF-OE^) or depleted (WM164^MITF-KD^), and the respective controls (C8161^EV^ and WM164^EV^). Scale bars: 200 μm. **(B,D)** p27^Kip1^ median fluoresecence intensity per cell plotted as a function of distance from the spheroid surface. C8161: n=13 (EV), n=14 (MITF-OE); WM164: n=5 (EV and MITF-KD); data: mean ±SEM; analysis: two-way ANOVA for the effect of MITF on p27^Kip1^ median fluorescence intensity per cell; **** p<0.0001.

These data suggest that MITF modulation of G1-arrest (Figure S 6A,B), and consequently phenotypic heterogeneity, are mediated through p27^Kip1^ which in turn is regulated by spheroid architecture.

### ROCK inhibition phenocopies the effects of high MITF levels on melanoma cells and spheroids

Dysregulation of Rho-ROCK-myosin signaling has been extensively linked to cancer (67–69). Recently, Rho-ROCK-myosin signaling has been shown to play a central role in melanoma acquired therapy resistance (70). As we have shown an inverse impact of MITF expression level on mechanosensitive signaling and cell contractility (Figure 5), we hypothesized that this phenomenon is driven by the Rho-ROCK-myosin pathway. To test this hypothesis, we studied the effect of ROCK inhibition on MITF^low^ cells and spheroids (C8161^EV^ and WM164^MITF-KD^) using the small molecule ROCK inhibitor Y27632, to assess whether reducing Rho-ROCK-myosin pathway activity could mimic and rescue the effect of MITF overexpression and depletion, respectively.

Indeed, similar to C8161^MITF-OE^ cells, ROCK-inhibited C8161^EV^ cells (EV+Y) contracted the collagen matrix significantly less than untreated C8161^EV^ cells, whereas ROCK inhibition of WM164*^MITF-KD^* cells (KD+Y) neutralized the increased matrix-contraction ability conferred by MITF depletion (Figure 7A). In spheroids, ROCK inhibition of C8161^EV^ and WM164*^MITF-KD^* also phenocopied and reverted, respectively, the cell morphology change caused by MITF overexpression, in particular, cell dendricity, which was prominent in untreated C8161^EV^ and WM164*^MITF-KD^* spheroids (Figure 7B). We then assessed spheroid solid stress, which was reduced to similar levels both by MITF overexpression and ROCK inhibition. Inhibition of ROCK in WM164^MITF-KD^ spheroids returned solid stress to a lower level comparable to that observed for WM164^EV^ spheroids (Figure 7C). By SPIM imaging we confirmed that, similar to C8161^MITF-OE^ spheroids, the equatorial diameter of ROCK-inhibited C8161^EV^ spheroids was increased, while the polar diameter was reduced, compared to C8161^EV^ (Figure 7D). Also in terms of p27^Kip1^ protein level ROCK inhibition mimicked the effect of MITF overexpression and rescued the effect of MITF knock-down (Figure 7E, Figure S 7A,B). We finally asked whether the effect of ROCK inhibition on solid stress was accompanied by changes in phenotypic heterogeneity as well, to support a mechanistic link between the two phenomena. ROCK inhibition, similar to MITF overexpression, significantly reduced the G_1_-arrested area in C8161^EV^ spheroids, and thus phenocopied the MITF-induced loss of phenotypic heterogeneity (Figure 7F, Figure S 7C). Similarly, the increase in phenotypic heterogeneity driven by MITF depletion in WM164 spheroids was reverted by ROCK inhibition, which led to a reduction of the G1-arrested area (Figure 7F, Figure S 7D).

**Figure 7:**
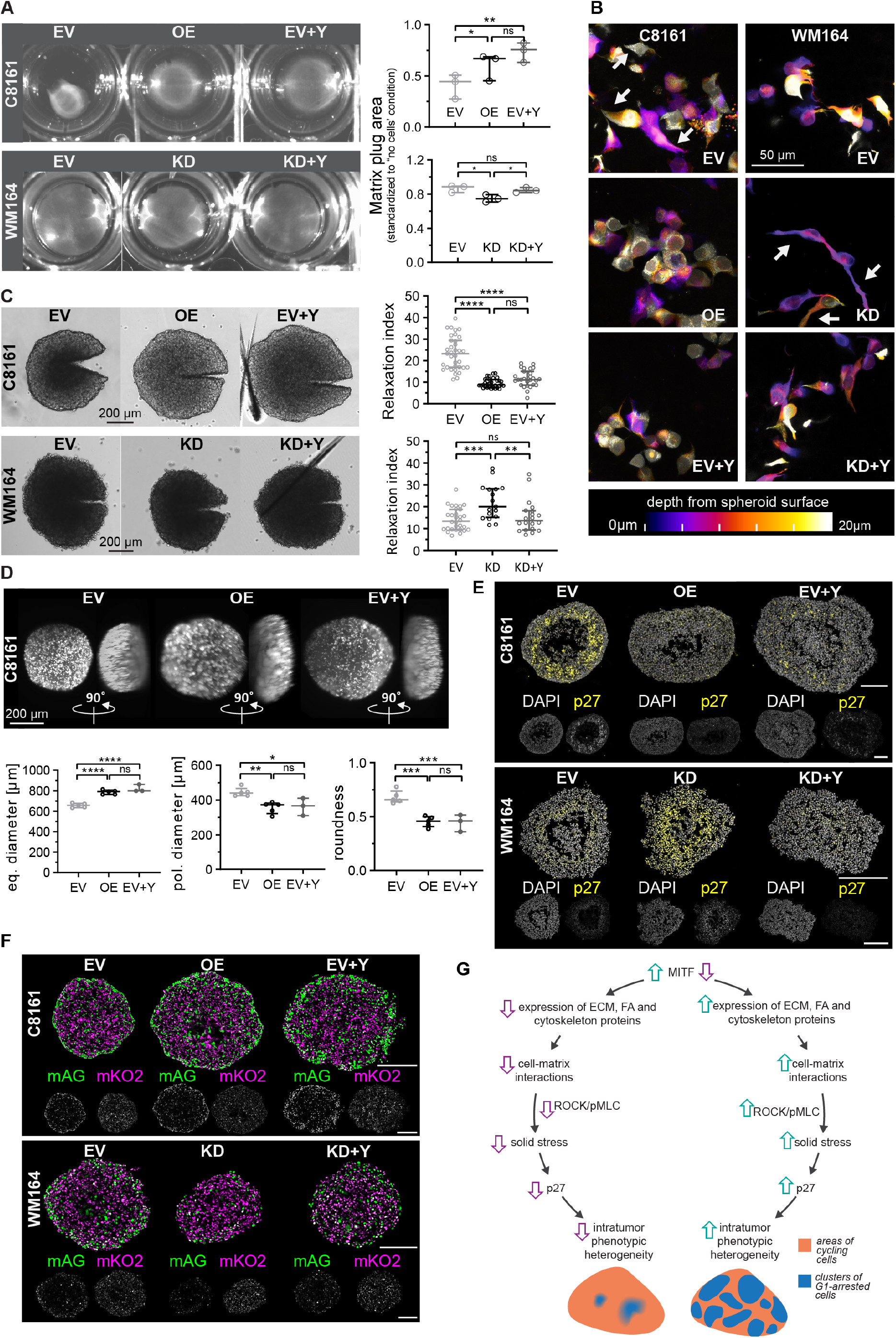
ROCK inhibition phenocopies the effects of high MITF levels on melanoma cells and spheroids. C8161^EV^, C8161^MITF-OE^, WM164^EV^ and WM164^MITF-KD^ cells were cultured with or without 10 μM ROCK inhibitor Y27632 (+Y). **(A)** Representative images of matrix contraction assay and quantitation of matrix plug contraction. **(B)** Representative depth color-coded confocal images of spheroid cells. Arrows indicate dendrites. **(C)** Representative transmitted-light microscopy images of incised spheroids and quantitation of their relaxation index. **(D)** Representative SPIM images (utilizing mAG for fluorescence) of spheroids from top view (equatorial) and side view (polar) and quantitation of equatorial and polar spheroid diameters and their ratio (roundness). Magenta and cyan lines indicate measured equatorial and polar diameters, respectively. **(E)** Representative p27^Kip1^ immunofluorescence of spheroid equatorial cryosections. **(F)** Representative mAG and mKO2 fluorescence images of spheroid equatorial cryosections. Scatter dot plot of data median and interquartile range; each dot represents the mean of an independent experiment (A,C) and a spheroid from three independent experiments (D); analysis: one-way ANOVA test with Tukey’s multiple comparisons test, ****p<0.0001, ***p<0.001, ** p<0.01, * p<0.05, ns p>0.05. **(G)** Schematic summary of the proposed mechanism of action..

Altogether, our findings show that ROCK inhibition phenocopies the effects of MITF overexpression on the melanoma cells’ ability to sense and respond to the microenvironment, leading to a decreased cell and ECM contraction. MITF overexpression and ROCK-inhibition both resulted in flatter spheroids with a lower solid stress and decreased phenotypic heterogeneity. This indicates that spheroid cell mechanotransduction, spheroid architecture and spheroid phenotypic heterogeneity are interrelated processes mediated by MITF through Rho-ROCK-myosin signaling.

## Discussion

Phenotypic heterogeneity is a major culprit of cancer therapy failure (3–6). We demonstrate that phenotypic heterogeneity is controlled through tumor cell-ECM crosstalk resulting in altered tumor microarchitecture, mechanotransduction and Rho-ROCK-myosin signaling. Melanoma shares these physical properties with any solid cancer underscoring the importance of our findings for therapeutically targeting this phenomenon.

In solid tumors, cancer cells crosstalk with their tumor microenvironment (TME) composed of the ECM and a variety of non-neoplastic cells. This process influences the survival, proliferation and dissemination of cancer cells (71). Spatial heterogeneity is an essential aspect of the TME, typically involving biochemical (e.g., oxygen concentration) and biomechanical (e.g., substrate stiffness) variations. TME spatial heterogeneity results in different cellular phenotypes throughout the tumor (72). Traditionally, clones of genetic mutations have been accepted as the cause of intratumoral phenotypic heterogeneity. The manifestation of distinct phenotypic traits by isogenic cells receiving different information from the environment is a more recent concept that has however already gained a fundamental role in intratumor heterogeneity (71,73). Undoubtedly, consideration of this aspect is crucial for a more accurate and complete understanding of cancer and its management.

Isogenic melanoma spheroids and xenografts are composed of differentially cycling tumor cells in a subcompartment-specific distribution, i.e. areas of proliferating and areas of G_1_-arrested cells (37), reflecting similar findings in patient samples (74). As this phenomenon is rapidly reversible in response to microenvironmental stress (37), we coined the term dynamic heterogeneity (75). MITF has been largely implicated in melanoma plasticity, yet its functional role is only partially elucidated and, at times, has been controversial. Most studies have either focused on the role of MITF in adherent cell culture, omitting the tumor 3D microenvironment, or *in vivo*, with limitations of deciphering detailed underlying mechanisms (41). Our 3D melanoma models allow us to establish an in-depth mechanistic understanding of this process, with a focus on important aspects provided by the 3D tumor environment, such as tumor architecture and solid stress. In this study, we demonstrate that MITF-mediated melanoma cell-ECM interaction alterations regulate phenotypic heterogeneity in melanoma via the Rho-ROCK-myosin signaling pathway.

The MITF rheostat model plays a complex role in melanoma cell proliferation through a number of direct effects on cell cycle proteins (17–19,21,76). However, MITF can also indirectly influence proliferation by affecting processes that regulate cell homeostasis, for instance metabolism, autophagy and survival (12). As opposed to direct effects on cell cycle proteins, the latter scenario relies on an adaptation to microenvironmental cues to manifest. This mechanism of action appears to apply to the control of phenotypic heterogeneity by MITF that we have identified in this work. Indeed, modulation of MITF expression did not impact the cell cycle profile in adherent culture, but only when the cells were surrounded by a G_1_-arrest promoting environment, as typically found in 3D models *in vitro* and *in vivo* and characterized by paucity of oxygen, high cell-cell contact and spatial confinement (77,78).

Insight into the interplay between MITF and the ECM is limited, mainly in the context of metalloproteinases and matrix degradation (79,80). A recent study on the effects of the tumor microenvironment on MITF expression demonstrated that a stiffer matrix leads to increased *MITF* mRNA levels in melanoma cell lines (81). Another study identified a correlation between progressive loss of extracellular matrix genes and *MITF* gene downregulation during tumor progression, which was modelled through pseudotime dynamics analysis of melanoma single-cell transcriptomes (82). We identified alteration of ECM organization upon modulation of MITF levels. ECM structure and organization are controlled by the nature and abundance of the ECM components as well as by the interaction between cells and the ECM via focal adhesions and cytoskeleton contractility. While we found that components belonging to both processes were affected upon MITF level modulation, the overlap of specific protein expression changes amongst cell lines was only partial. These findings are explained by a phenomenon typically observed in cancer, especially in cancers with high mutational burden such as melanoma, where the same cell adaptation, in this case alteration of the ECM organization, is achieved via diverse cell signaling remodeling (83). Not surprisingly, we did not find a distinct common molecular signature, but rather a shared alteration in three key processes involved in in the regulation of tumor architecture, which together drive tumor growth and therapy response: ECM composition, cell-matrix adhesion and cell cytoskeleton functionality. These three aspects are pivotal for the inside-out/outside-in signaling that mediates reciprocal regulation between the cell and the ECM.

Inside-out/outside-in signaling is mediated by actin stress fibers, rich in myosin II motors, which attach to focal adhesion complexes and transmit forces from the surrounding ECM to the cells and *vice versa* (84,85). This mode of signaling allows for remodeling, organization and contraction, of the surrounding ECM. An enhanced inside-out/outside-in signaling is therefore consistent with the increased elongation of focal adhesions, the cells’ ability to contract the surrounding matrix and the ECM organization that we observed upon MITF depletion. MITF knockdown led to increased focal adhesion numbers accompanied by raised levels of the MITF direct target paxillin in adherent cultures (86). We found that focal adhesions did not change in number upon MITF depletion in spheroids, however they had a more elongated shape. This phenomenon was accompanied by increased protein levels of zyxin, fascin and integrin-α4, while paxillin did not change (proteomics analysis, data not shown). Zyxin, fascin and integrin-α4 are focal adhesion components, whose genes are bound by MITF based on ChIP-seq analysis. Moreover, all three correlated negatively with MITF in expression datasets from three independent panels of melanoma cell lines in adherent culture and in clinical samples. This indicates that MITF controls focal adhesion maturation by directly regulating the expression of zyxin, fascin and integrin-α4.

Actomyosin contractility is also pivotal to focal adhesion maturation and functionality. Contractile force is mediated by the Rho-ROCK-myosin signaling pathway, whose inhibition we found to phenocopy the phenotype caused by MITF overexpression, both at the cellular and spheroid level. Taken together, our findings imply an inverse link between the Rho-ROCK signaling axis and MITF levels, which is in line with increased Rho activity, ROCK-dependent invasiveness and increased myosin phosphorylation found in melanoma cells where MITF was depleted (18,79,87).

There is a growing body of evidence on the impact of mechanical cues of the microenvironment on cancer cells biology, including cancer cell plasticity (88). Extensive investigation has been carried out on the effects of ECM stiffness on tumorigenesis, and the contribution of solid stress to tumor biology is an exciting emerging field of research (46,89,90). In our study, in addition to decreased phenotypic heterogeneity, high MITF expression levels drove two main phenotypes: morphological relaxation and decreased solid stress. It has been shown that compressive stress in cancer cell spheroids led to ROCK-dependent G_1_-arrest via p27^Kip1^ upregulation, which in turn was triggered by cell volume reduction (66). Our results are in agreement with these findings, whereby solid stress and phenotypic heterogeneity changed simultaneously in our spheroids when MITF expression was altered and were dependent on ROCK activation.

In melanoma, MITF was shown to repress the expression of the *DIAPH1* gene encoding the diaphanous-related formin 1 (Dia1) that promotes actin polymerization and to increase p27^Kip1^ protein level leading to cell cycle G1-arrest (18). While we observed changes in p27^Kip1^ level in adherent cultures upon MITF expression modulation (Figure S 6), these were not accompanied by variations of cell cycle distribution (Figure S 1G). Furthermore, our proteomics analysis indicated that Dia1 protein levels were not affected by MITF overexpression or knockdown (data not shown). Elevated activity of the Rho-ROCK-myosin signaling axis is typically associated with increased proliferation via promotion of cell contractility, cell adhesion, cytokinesis, oncogene activation and regulation of cell cycle proteins. Nevertheless, there is a number of studies reporting an opposite role of the Rho-ROCK-myosin signaling pathway through negative feedback on growth factor signaling (91) and promotion of tumor suppressor genes such as PTEN (92,93), p21 (94,95) and p27^Kip1^ (18,96,97).

We propose that MITF controls ECM organization and mechanical properties of melanoma through regulation of cell-mediated matrix contraction. By acting as a scaffold and a capsule, the ECM in turn controls spheroid solid stress and physical pressure undergone by cells, regulating their ROCK-mediated contractility and ultimately their proliferation.

While MITF is mainly dysregulated in melanoma, ECM and ROCK signalling alterations are common to other solid cancers and, importantly, influence cancer cell plasticity, disease progression and therapy response (34,35,69,88,98,99). It is therefore plausible that a similar ECM-cell interaction mechanism controlling tumor heterogeneity occurs in other solid cancers. In fact, wide scientific interest and growing efforts are directed to the development of anti-cancer therapy targeting the crosstalk between the ECM and tumor cells (100,101). This work demonstrates a novel role of MITF in melanoma phenotypic heterogeneity and underscores the ECM-cancer cell interaction as a potential therapeutic target not only for melanoma, but also for other solid tumors.

## Methods

### Cell lines

1205Lu-C5, 451Lu-Z1, C8161-A7, WM164-F11, WM793-G10, WM983B-X2, WM983C-Y3 were generated at the Centenary Institute (for a review of correlation functions see 37) and were made available at the University of Queensland after approved material transfer agreements from all involved parties. All cell lines were genotypically characterized (22,103–105) and authenticated by STR fingerprinting (Analytical Facility, QIMR Berghofer Medical Research Institute, Herston, Australia). None of the cell lines contain the MITF E318K mutation; WM793 and 1205Lu carry the synonymous E388E mutation (pers. comm. Katherine Nathanson, University of Pennsylvania, Philadelphia, PA, USA).

### Mice

Experiments were approved by the University of Sydney Animal Ethics Committee in accordance with the guidelines from the National Health and Medical Research Council (Ethics #K75/10-2008/3/4910). CB17 NOD/SCID mice were provided by the Animal Resources Centre, Canning Vale, Australia, and housed in the Centenary Institute Animal Facility, Newtown, Australia.

### Plasmids

The plasmids generated in this paper were made at the University of Queensland and are available after approved material transfer agreement.

### Cell culture

All Melanoma-FUCCI cells were cultured in melanoma cell medium: 80% MCDB-153 medium, 20% L-15 medium, 4% fetal bovine serum, 5 μg/mL insulin and 1.68 mM CaCl2 as per methods published (40).

### Generation of MITF knockdown cell lines

The lentivirus was produced by co-transfection of human embryonic kidney 293T cells with four plasmids, including a packaging defective helper construct (pMDLg/pRRE), a Rev plasmid (pRSV-Rev), a plasmid coding for a heterologous (pCMV-VSV-G) envelope protein, and MISSION® pLKO.1-puro Empty Vector Control Plasmid DNA (PLKO), MISSION® pLKO.1-puro Non-Mammalian shRNA Control Plasmid DNA (Scram). The FUCCI-expressing single cell clones WM164-F11, WM983B-X1 and WM983C-Y3 were transduced with MITF MISSION® shRNA Lentiviral Transduction Particles, PLKO or Scram lentivirus, per manufacturer’s directions. Transfected cells were selected and cultured in melanoma cell medium containing 1 µg/ml puromycin. To allow for better visibility for color-deficient readers, monomeric Kusabira Orange2 (mKO2; G1 phase) and monomeric Azami Green (mAG; S/G2/M) fluorescence is shown in magenta and green, respectively.

### Generation of MITF overexpression cell lines

The open reading frame for the MITF-M was amplified from cDNA generated from A2058 cell line using the primers:

> MITF-LVX-Eco-F: 5’-GCGCGAATTCACCATGCTGGAAATGCTAGAA-3’
>
> MITF-LVX-Eco-R: 5’-GCGCGAATTCCTAACAAGTGTGCTCCGT-3’

The primers contained *Eco*RI restriction sites before the start codon and after the stop codon as well as ACC nucleotides to create an optimal kozac sequence immediately 5’ to the ATG codon. Insert was cloned into the *EcoRI* site of the pLVX-PURO vector and the orientation and sequence of the full open reading frame confirmed using pLVX forward and reverse primers. Lentivirus with pLVX-PURO-MITF-M (MITF OE) and with pLVX-PURO vector only (EV) plasmids were generated using Lenti X-HTX packaging system as per manufacturer’s instructions.

The FUCCI-expressing single cell clones C8161-A7, WM793b-G10 were transduced with MITF-OE or EV lentivirus in 4 µg/ml polybrene. Transfected cells were selected and cultured in melanoma cell medium containing 1 µg/ml puromycin.

Cells were routinely tested for mycoplasma using PCR (106).

### Xenograft studies

The flanks of 6-week-old female CB17 NOD/SCID mice were injected subcutaneously with 2×10^6^ FUCCI-transduced melanoma cells in 100 µl Melanoma cell medium as previously described (107). Mice were weighed thrice weekly and tumor growth measured with digital calipers. Age-matched or size-matched tumors were harvested and compared. Mice were sacrificed when tumors reached 1 cm^3^ volume or became ulcerated.

Following sacrifice, mice were perfusion fixed by cutting the inferior *Vena cava* and injecting 10 ml PBS, followed by 8 ml 4% formaldehyde into the left ventricle of the heart. Tumors were surgically excised and placed in 4% formaldehyde for 18-24 h at RT. Tumors were cut into 200-μm sections using a vibratome. Sections were placed into ice cold 70% ethanol for a minimum of 1 h at 4°C. For imaging, sections were mounted on glass slides in anti-fade media: 0.25% (w/v) DABCO and 90% (v/v) Glycerol 90 ml in PBS) with cover slip held in place using vacuum grease.

### Tumor cell clustering image analysis

Z-stacks of tumor sections were collapsed into an extended focus image. Red and green nucleus masks were created using Volocity software. The coordinates of cell centroids were then exported for the clustering analysis (Figure 1C, F). Tumors with large areas of necrosis (no nuclei) were excluded from the analysis.

We adapted the astrophysical correlation function to measure the degree of cell co-location or clustering within a tumor (for a review of correlation functions see 108). The correlation function measures the excess (or diminished) probability, dP, of finding a cell at a distance, r, from another cell within some region, dA. It is defined as, dP = (1 + ξ(r)) N dA / A, where N is the number of cells within the area, A.

To explicitly account for the effects of the tumor boundary, the correlation function of green and red cells were measured using the Szalay & Landy estimator (39):

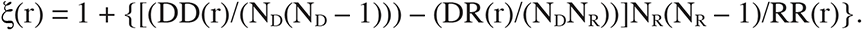

It works by comparing the distribution of cell separations to a completely random sample generated by us. DD(r), RR(r) and DR(r) are the number of unique cell-cell, random-random and cell-random pairs at a separation of r. The pair counts were normalized by the total number of cell-cell, random-random and cell-random pairs. Random samples were generated with 2-80x density compared to the number of cells in each sample, to ensure that the random sample density is high enough to measure the effect of the tumor boundary on the correlation function. To diminish the effect of an individual random sample on our correlation function measurements, five random samples were generated for each cell sample, and the results were averaged.

The final clustering difference plots were generated in two steps. First, for each tumor image the difference in red and green cell clustering were converted into an absolute difference and then the mean absolute difference for each set of tumor images were calculated. Finally, the mean absolute difference plots were scaled by the standard error of the mean for each tumor set. P-values of 0.05, 0.01, 0.005, 0.001 and 0.0001 correspond to sigma values of 1.96, 2.58, 3.81, 3.29 and 3.89, in the sigma difference plots.

### Spheroid formation

All spheroids were produced by method described previously (40). For ROCK inhibitor treatment, spheroids were treated with 10 µM Y-27632 at the time of spheroid formation. On day 4 of the spheroid formation, spheroids were collected for immunofluorescence, immunoblots or other following assays.

### Spheroid cryosectioning

As described previously (42), day 4 spheroids were fixed in 4% paraformaldehyde solution at room temperature for 20 minutes and washed thoroughly in phosphate buffer saline (PBS). The spheroids were then embedded in tissue freezing medium and frozen on dry ice. The molds were then sectioned at 20µm thickness using ThermoFisher HM525NX Cryostat.

### Immunostaining

#### Spheroid section immunostaining

The spheroid sections on glass slides were permeabilized using 0.1% Triton and blocked with 5% BSA-PBS or ABDIL buffer (Cold Spring Harbor Protocols). The sections were then stained with anti-pimonidazole (1:200, Hypoxyprobe) and the secondary antibody anti-Rabbit IgG-647 (1:1000) in respective blocking buffer with adequate washing with PBS between antibody additions. The sections were stained with 4′,6-diamidino-2-phenylindole (DAPI) before being mounted using Mowiol mounting media (Cold Spring Harbor Protocols). The samples were imaged using Laser Scanning Confocal Microscope Olympus FV3000: UPlanSApo 10X – NA 0.4, UPlanSApo 20X – NA 075.

#### 2D immunostaining

Cell grown on No. 1.5 coverslips were fixed with 1X BRB80 (Cold Spring Harbor Protocols) containing 4% paraformaldehyde solution at 37°C for 12 min, washed in 1xPBS, permeabilized using 0.1% Triton for 5 min and blocked with ABDIL buffer. The primary antibodies: anti-pFAK (1:200) and anti-pMLC (1:50) and the secondary antibodies: anti-Rabbit IgG-647 (1:1000), anti-Rabbit IgG-750 (1:1000) were added in ABDIL buffer with adequate washing with TBS-T (0.1% Tween) between antibody addition. The samples stained with DAPI, mounted using Mowiol mounting media and imaged using Laser Scanning Confocal Microscope Olympus FV1200: UPlanSApo 60X Oil – NA 1.35. Z-stack images of 1µm thickness were obtained and individual channels were combined to a single image by maximum intensity projection using ImageJ software.

### CellProfiler analyses

#### Percentage of mAG^+^ cells

Spheroids were identified as primary objects using a binary image obtained from the merged mAG and mKO2 images. mAG^+^ and mKO2^+^ cells were identified as primary objects based on their size and their mAG and mKO2 signal intensity, then their distance from the spheroid surface measured (minimum child-parent distance). mAG^+^ and mKO2^+^ cells were counted in bins of distances from the spheroid surfaces (10μm increments) and the percentage of mAg^+^ cells was calculated for each bin.

% of mAG^+^ cells= mAG^+^ cells/ (mAG^+^ cells + mKO2^+^ cells)

Figure 2: n = 16 (WM164^EV^), n =16 (WM164^MITF-KD^), n = 11 (C8161^EV^), n = 12 (C8161^MITF-OE^).

Figure 2S: n = 7 Figure 2S: (WM983B^EV^), n = 7 (WM983B^MITF-KD^), n = 8 (WM793B^EV^), n = 7 (WM793B^MITF-OE^).

Figure 8: n = 6 (C8161^EV^), n = 6 (C8161^MITF-OE^), n = 6 (C8161^EV + Y^), n = 6 (WM164^EV^), n = 6 (WM164^MITF-KD^), n = 9 (WM164^MITF-KD + Y^).

Data: mean ±SEM; analysis: two-way ANOVA for the effect of MITF on % of mAg2^+^ cells

#### Mean intensity of pMLC2

DAPI images were used to identify cell nuclei as primary objects while secondary objects were identified based on pMLC2 images. Mean intensity pMLC2 signal was measured for each cell based on identified secondary objects; n = 58 (C8161^EV^), n = 62 (C8161^MITF-OE^), n = 60 (WM164^EV^), n = 73 (WM164^MITF-KD^) cells from 3 independent experiments; analysis: Mann Whitney test.

#### Focal adhesion eccentricity

Focal adhesions were identified as primary objects based on their size and pFAK signal, then their count and shape eccentricity (referred to as elongation) were calculated. Considering each focal adhesion an ellipse, the eccentricity is the ratio of the distance between the foci of the ellipse and its major axis length. The value is between 0 and 1 (0 = circle, 1= line segment); n = 44 (C8161^EV^), n = 53 (C8161^MITF-OE^), n = 62 (WM164^EV^), n = 64 (WM164^MITF-KD^) cells from 3 independent experiments; analysis: unpaired t-test.

#### Elongated focal adhesion count

The number of properly formed elongated focal adhesions per cell were manually counted. Phosphorylated FAK signal that had defined edges were included in the count. n = 48 (C8161^EV^), n = 53 (C8161^MITF-OE^), n = 59 (WM164^EV^), n = 65 (WM164^MITF-KD^) cells from 3 independent experiments; analysis: Mann Whitney test.

### SPIM

Day4 spheroids were mounted in imaging chamber (109) using 1.5% low melting agarose and submerged in PBS. The submerged spheroids were then imaged using a diffused digitally scanned light-sheet microscope (110) with two light-sheets illuminating the sample. The scanning light sheet was generated using a 488nm laser (OBIS 488 lx), attenuated with a ND filter (Thorlabs NE10A), scanned with 2D galvo mirrors (Thorlabs GVSM002/M), a 50/50 beamsplitter, and a 1D line diffuser (RPC Photonics EDL-20-07337). One galvo scanning direction (Y) created the light sheet while the second direction (Z) created the depth scan in the sample. The two mirrors were driven independently using Arduinos (DUE) with custom-written code. The Y scanning was a sawtooth scan at 600 Hz, which was synchronized to the camera acquisition to ensure similar illumination for each camera acquisition. The Z galvo was driven in steps to scan the light-sheet through the sample. The 50/50 beamsplitter created two light sheets projecting orthogonally in the sample. Excitation was gathered by an Olympus Plan N 10x 0.25 NA objective and projected onto the PCO edge 5.5 camera with a combination of filter (F, Thorlabs FF01-517/520-25), tube lens (L3, 180 mm focal length, Thorlabs AC508-180-A), relay lenses (Lr, Thorlabs AC254-125-A-ML), ETL (Optotune EL-10-30-Ci-VIS-LD driven with Gardasoft TR-CL180) and offset lens (Lo, Eksma Optics 112-0127E). SPIM: objective information, section thickness, any post processing The imaging system was controlled using μManager, based on ImageJ (110,111). The exposure time was set at 100ms for each plane, with a laser power output of 40mW, which was attenuated to 1mW for each plane at the sample. The depth of scanning was adapted to each spheroid with 3 μm steps between each plane.

In our experiments, an exposure time of 10ms was chosen for each plane during volumetric imaging, with laser power output of 40 mW, which was attenuated to 1.5 mW for each plane at the sample.

Three-dimensional image reconstruction and the equatorial and polar diameter measurements were done using imageJ software. Spheroid roundness was calculated by polar diameter/ equatorial diameter.

### Immunoblot

Cells from 80% confluent flask or 4 day old spheroids were lysed in RIPA lysis buffer: 150mM NaCl, 5mM EDTA (pH 8.0), 50mM Tris (pH 8.8), 1% (v/v) NP-40 (IGEPAL CA-630), 0.5% (w/v) Sodium deoxycholate and 0.1% (w/v) SDS, quantified using Pierce™ BCA Protein Assay Kit and boiled in sample buffer: 30% (v/v) glycerol, 60mM Tris, 2% (w/v) SDS, 10% (v/v) β-mercaptoethanol, 0.1% (w/v) bromophenol blue. Alternatively, the cells were lysed directly in sample buffer containing 50mM Tris (pH6.8), 3% (w/v) SDS, 10% (v/v) glycerol, 100mM DTT and 0.1% (w/v) bromophenol blue and sonicated. All buffers contained 1x protease inhibitor cocktail, 1mM phenylmethylsulfonyl fluoride, 1mM sodium orthovanadate, 1mM Sodium fluoride, 1 ug/ml Aprotinin and 1ug/ml Leupeptin at the time of lysis and were boiled for 5 min at 95°C prior to use. Approximately 20-30µg protein was loaded per lane on NuPAGE™ 4-12% Bis-Tris Protein Gels and electro-blotted on to Immobilon-P PVDF Membrane [FIG1] or on 10% Bis-Tris gel and electro-blotted on to Immun-Blot® Low Fluorescence PVDF membrane [FIG2]. Membranes were blocked in 5% skim milk in TBS-T (0.1% tween) and probed with antibodies against: MITF (1:500), Rab27a (1:1000), GAPDH (1:5000). Following washes with TBS-T, bound antibodies were detected with anti-mouse IgG-HRP or anti-rabbit IgG-HRP (1:500) and Clarity™ Western ECL Substrate. The blots were then imaged using Bio-Rad ChemiDoc MP. The intensity of the bands was measured using imageJ and normalized before fold change difference was calculated.

### Flow cytometry for FUCCI

Cells were seeded on tissue culture plastic in melanoma cell medium at 40,000 cells/cm^2^ for 24 hours, dissociated using trypsin and analyzed using Flow Analyser LSR Fortessa X20. The number of mKO^+^ and mAG^+^ cells were calculated based on histogram and the %mKO^+^ cells were calculated.

% of mKO2^+^ cells= mKO2^+^ cells/ (mKO2^+^ cells + mAG^+^ cells)

N=3 independent experiments; analysis: unpaired t-test on % of mKO2^+^ cells.

### Proteomics

Cells were seeded on tissue culture plastic in melanoma cell medium at 40,000 cells/cm^2^ for 24 hours (2D) or 4-day old spheroids (3D) were used. The samples were washed in thoroughly in PBS, lysed in buffer containing 1% Sodium deoxycholate, 10mM tris(2-carboxyethyl)phosphine (TCEP), 40mM 2-chloroacetamide, 1x Protease inhibitor cocktail in 100mM Tris(pH 8) and sonicated at 4°C for 15 min in a water bath sonicator. The samples were clarified by centrifugation at 13000 x g, 4°C for 10 min. The supernatant was collected and the protein concentration was measured using direct detect. Protein samples of 10ug were denatured at 1ug/ul by boiling for 5 min at 95°C. The samples were diluted 1 in 10 with water before overnight digest with 0.2ug of trypsin at 37°C. Samples were acidified with 0.5% (v/v) trifluoro acetic acid and centrifuged at 13000 x g for 10 min. The supernatant was purified using Glygen C18 tips before being analyzed on Thermo Scientific Easy nLC 1000 with Q Exactive Plus orbitrap mass spectrometer. Peptides were loaded onto a 2mm x75um c18 trap column and separated with an Easy LC C18 analytical column 50cm x 50um over 100mins from 3% to 25% acetonitrile in 0.1% formic acid, followed by 40mins from 25% to 40% acetonitrile. Mass spectrum was acquired at 70K resolution, 350-1400m/z, 3e6 AGC with a maximum injection time of 100ms and data dependent ms2 was acquired at 17.5K resolution, 5e5 AGC maximum injection time of 55ms, TopN of 20, dynamic exclusion of 30s and exclusion of unassigned, 1, 8 and >8z. Mass spectrum data was searched against from Swiss-Prot – human species protein database (April 2017) using Spectrum Mill and Proteome Discoverer search engines with standard setting of tryptic peptides with a maximum of 2 miscleavages, fixed carbamidomethylated cysteine and variable oxidation of methionine. Only proteins with FDR of <1% were use in subsequent analysis. Filtering was performed for proteins reliably detected in 3 out of 4 technical replicates of at least one of the experimental groups. The analysis was performed by Queensland Cyber Infrastructure Foundation Ltd, The University of Queensland - St Lucia, QLD-4072. After filtering 2657 proteins remained. Differential proteins were determined by Analysis of variance (ANOVA) with correction for multiple testing uses the False Discovery Rate method from Benjamini and Hochberg. All analyses were carried out using the R statistical software (112).

Functional enrichment analysis of the differentially expressed proteins was performed using the STRING biological database and web resource based on the Genet Ontology (GO) functional classification system with default settings on 18^th^ of May 2020.

### Individual cell imaging in spheroids

Cells were stained with 10 µM CellTracker™ Deep Red Dye as per manufacturer’s protocol and mixed with unstained cells at 1:50 ratio respectively before spheroid formation. The spheroids were treated with 10 µM Y-27632 every day for treatment condition. On day 4, spheroids were fixed and cleared using CUBIC based method as described previously (113). The spheroids were incubated in 50% (v/v) CUBIC Reagent 1A at room temperature for 1 h and then in 100% CUBIC reagent 1A at 37°C for 6-8 h till the spheroids were transparent. They were then mounted in CUBIC Reagent 2 overnight for refractive index matching. Images were acquired using Laser Scanning Confocal Microscope Olympus FV3000: UPlanFL N 40X Oil – NA 1.3. Z-stack images of 1µm thickness were obtained and images of CellTracker™ stained cells were temporally color coded and stacked for observation using ImageJ software.

### Accumulated solid stress analysis in spheroids

Manual incisions were made using an 11 blade on day 4 spheroids ranging between 35-55% of the spheroid diameter and imaged using Inverted Manual Microscope Olympus IX73: UPlanFL N 10x – NA 0.30. The spheroid diameter (d), incision depth (i), and the opening distance (a) were measured using ImageJ software. The relaxation indices (RI) were calculated using the formula: RI = a/ d *100. Experiments were repeated three times with multiple technical replicates and statistical significance calculated using the Mann Whitney test or one-way ANOVA test with Tukey’s multiple comparisons test in GraphPad Prism.

### Collagen contraction assay

Collagen solution of 2mg/ml was made using 40% (v/v) bovine type 1 collagen, 10% (v/v) 10x EMEM, 1% (v/v) GlutaMAX™, 2% (v/v) Sodium bicarbonate and 10% (v/v) fetal bovine serum. The working stock of cell suspension was made in melanoma cell medium at 2×10^5^ cells/ml. The two solutions were mixed 1:1 and 500µl of solution was added per well of 24 well plate and incubated at 37°C immediately for 30 minutes. Once the gel solidified, 1.5ml medium per well was added with or without 10 µM Y-27632. The collagen plug was gently released from the edges and base using 200 µl pipette tip and the plates were incubated at 37°C. The plates were imaged on day 4 using Bio-Rad ChemiDoc MP to obtain a quick snapshot of the whole plate and the area of the plugs were measured using ImageJ. The percentage of contraction was calculated in relation to acellular plugs.

Matrix plug area = area of cell treated collagen plug / area of acellular collagen plug Experiments were repeated three times with three technical replicates for statistical significance and analyzed using paired t-test or ANOVA with test with Tukey’s multiple comparisons test.

### Gene expression profile and ChIP-seq analysis

In this study, the expression data of melanoma of selected genes were retrieved and downloaded from the CCLE (https://portals.broadinstitute.org/ccle), TCGA (https://portal.gdc.cancer.gov/), Tsoi et al (114) and Goding lab. Heatmaps were generated from normalised gene expression data using the Pheatmap package in R (https://cran.r-project.org/web/packages/pheatmap/index.html). UCSC browser screenshots for duplicate MITF ChIP experiments were generated from the dataset deposited in the Gene Expression Omnibus under accession number GSE77437 generated by (115). The Raw fastq files were processed and mapped to the human genome build hg19 (GRCh37, February 2009) allowing for 2 mismatches using Bowtie 1.1.2 (102). Duplicate reads were eliminated using PicardTools version 1.96, http://picard.sourceforge.net and the University of Santa Cruz (UCSC) Genome Browser was used to examine library-count normalized read density at gene loci of interest.

## Acknowledgments

This research was carried out at the Translational Research Institute (TRI), Woolloongabba, QLD, and at the Centenary Institute (CI), Camperdown, NSW, Australia. TRI is supported by a grant from the Australian Government. We thank the staff in the core facilities at TRI and CI for their outstanding technical support (microscopy, flow cytometry, proteomics, histology and animal facility). We thank Prof. Atsushi Miyawaki, RIKEN, Wako-city, Japan, for providing the FUCCI constructs, Prof. Meenhard Herlyn and Ms. Patricia Brafford, The Wistar Institute, Philadelphia, PA, for providing the cell lines, Dr. Miklós Geiszt for providing the peroxidasin antibody, Ms. Dorothy Loo-Oey, TRI, for the proteomics study, and Prof. Jochen Guck, Max-Planck Institute for the Science of Light, Erlangen, and Dr. Anna Taubenberger, Technische Universität Dresden, Germany, for their intellectual input. N.K.H. is a Cameron fellow of the Melanoma and Skin Cancer Research Institute, Australia. K.A.B. is a fellow of Cancer Institute New South Wales (13/ECF/1-39). D.S.H. is a David Sainsbury Fellow of the National Centre for the Replacement, Refinement and Reduction of Animals in Research. This work was supported by project grants to N.K.H.: APP1003637 and APP1084893 (National Health and Medical Research Council); RG 09-08 and RG 13-06 (Cancer Council New South Wales), 570778 and 1051996 (Priority-driven collaborative cancer research scheme/Cancer Australia/Cure Cancer Australia Foundation), 08/RFG/1-27 (Cancer Institute New South Wales), and Meehan Project Grant 021174 2017002565. JC and CRG are supported by the Ludwig Institute for Cancer Research.

## Author Contributions

Conceptualization, Supervision, Project Administration and Funding Acquisition, N.K.H.; Intellectual Contribution to Conceptualization and Funding Acquisition: W.W., H.S., B.G.; Methodology, N.K.H., E.K.S., C.R.G.; Investigation, L.S., C.A.T.-M., G.P.G., D.S.H., K.A.B., R.J.J., G.C.V., M.E.F., S.M.D.-M., N.K.H.; Validation and Formal Analysis, L.S., C.A.T.-M., G.P.G., D.S.H., K.A.B., R.J.J., J.C., N.M., G.M.B., C.R.G., N.K.H.; Data Curation: L.S., N.K.H.; Writing – Original Draft, L.S., N.K.H.; Writing – Review & Editing, L.S., S.M.D.-M., A.G.S., S.J.S., H.S., B.G., W.W., C.R.G., N.K.H.

---

**Figure S 1:**
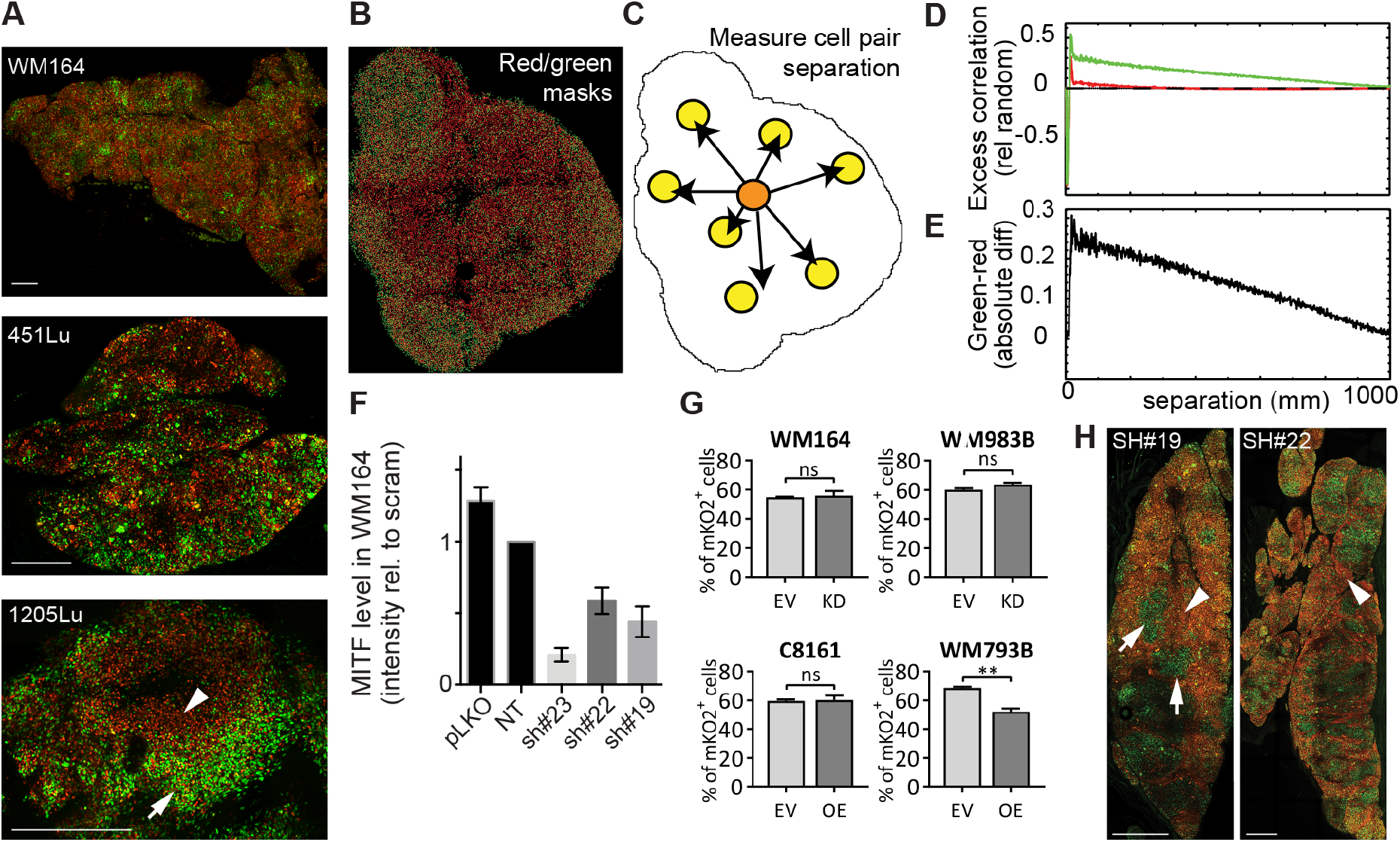
Phenotypic heterogeneity in melanoma xenograft tumors in vivo is dependent on expression levels of MITF, clustering analysis method, MITF expression levels in WM164 cells and percentage of mKO2^+^ cells in adherent cultures. **(A)** Representative examples of predominantly homogeneously proliferating MITF^high^ (WM164 and 451Lu) and clustered MITF^low^ (1205Lu) xenografts. Arrows and arrowheads indicate proliferating and G_1_-arrested clusters, respectively; scale bars: 0.5 mm. **(B)** Clustering analysis method using Figure 1B (low MITF, C8161) as an example. Red/green nuclei masks (centroids) automatically generated in Volocity. Note that not all cells could be separated due to the high density – but the overall red/green cell pattern matches the original image (Figure 1B). **(C)** The distance of each green cell to every other green cell in the image was measured; the same process was repeated for the red cells. **(D)** A random sample was generated within the tumor border. Histograms of red-red, red-random, green-green, green-random and random-random pair separations were combined to measure the excess/diminished red-red and green-green pairs at a given separation, while taking into account the tumor border. Note that for this image there is an accumulation of green cell pairs (relative to the random sample – dotted line) separated by shorter distances. This indicates that the green cells are showing increased clustering at these distances. **(E)** To combine the analyses of multiple tumors, we used the mean of the absolute differences between the green and red excess correlation curves. We identified differences in red and green clustering irrespective of which was stronger. Here we show the absolute difference curve for this particular tumor. **(F)** MITF expression level measured by Western Blot of WM164 cells expressing vector only (PLKO), scrambled shRNA (scram) and three different shRNA targeting MITF (sh#23, sh#22 and sh#19). **(G)** Representative examples of clustered xenografts generated with MITF-depleted cells (sh#19 and sh#22). Arrows and arrowheads indicate proliferating and G_1_-arrested clusters, respectively; scale bars: 1 mm. **(H)** Percentage of mKO2^+^ cells (mKO2^+^ cells / (mKO2^+^ cells + mAG^+^ cells)) in adherent cultures of WM164, WM983B, C8161 and WM793B; parental (PAR), SCR (scrambled shRNA), EV (empty vector), KD (MITF knock down) and OE (MITF overexpression).

**Figure S 2:**
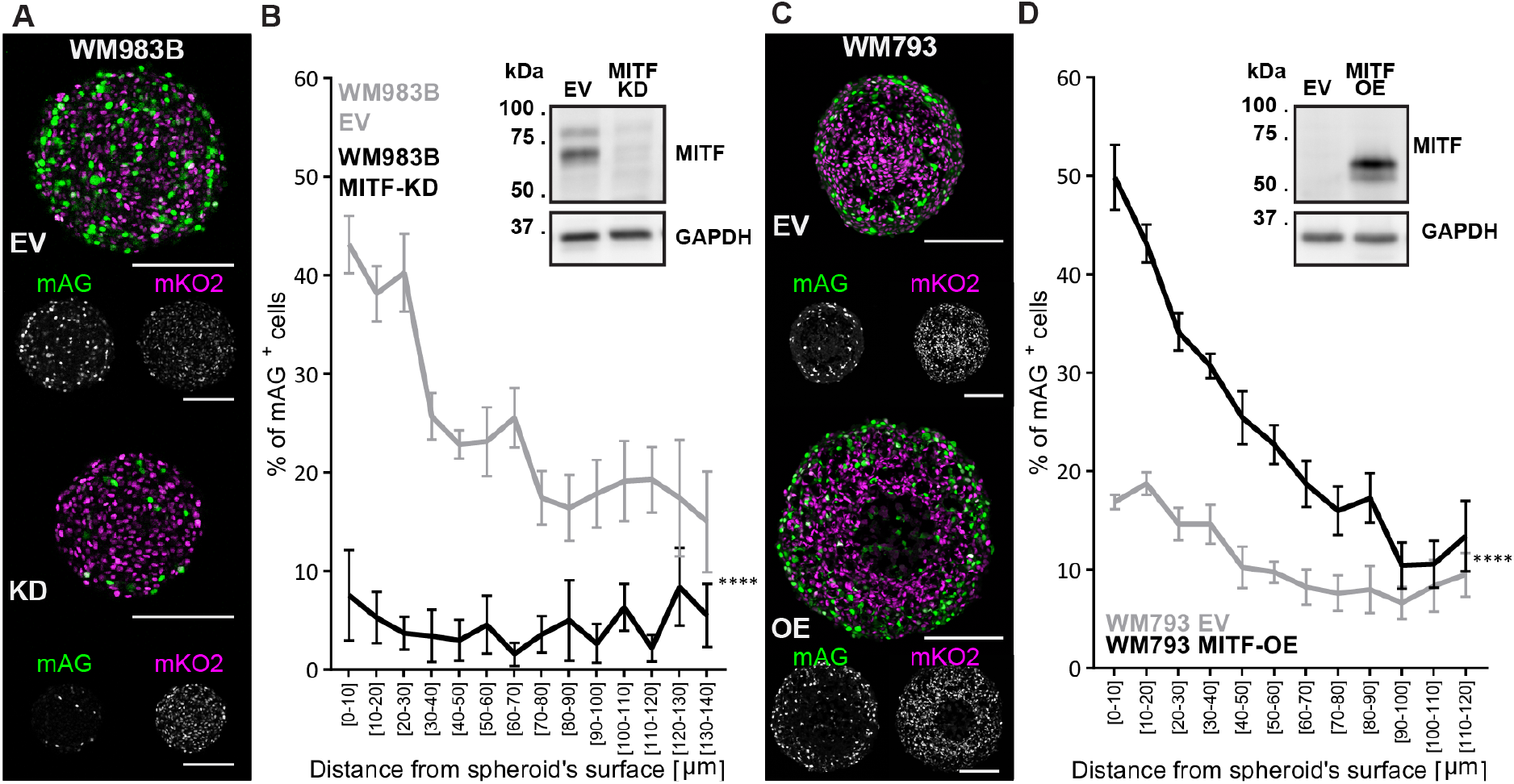
Phenotypic heterogeneity in melanoma xenograft tumors in vitro is dependent on expression levels of MITF. **(A,C)** Representative FUCCI fluorescence images (monomeric Azami green, mAG, green; monomeric Kusabira Orange2, mKO2, magenta) of equatorial cryosections of spheroids with MITF depleted (WM983B^MITF-KD^) or overexpressed (WM793B^MITF-OE^), and the respective controls (WM983B^EV^ and WM793B^EV^). Areas of mAG^+^ and mKO2^+^ cells proliferate, while zones enriched with mKO2^+^ cells are G_1_-arrested; scale bars: 200 μm. **(B,D)** Analysis of spatial distribution of proliferation. Graphs show the percentage of mAG^+^ cells (mAG^+^ cells/(mAG^+^ cells+mKO2^+^ cells)) as a function of distance from the spheroid surface. Note, cells beyond specific spheroid depths (cell-line dependent) were omitted from the analysis because of high variability driven by core necrosis. WM983B: n=7 (EV), n=7 (MITF-KD); WM793B: n=8 (EV), n=7 (MITF-OE); data: mean ±SEM; analysis: two-way ANOVA for the effect of MITF on % of mAG^+^ cells; **** p<0.0001. Immunoblots show the level of MITF knockdown (B) and overexpression (D). EV, empty vector; KD, knock-down; OE, overexpression.

**Figure S 3:**
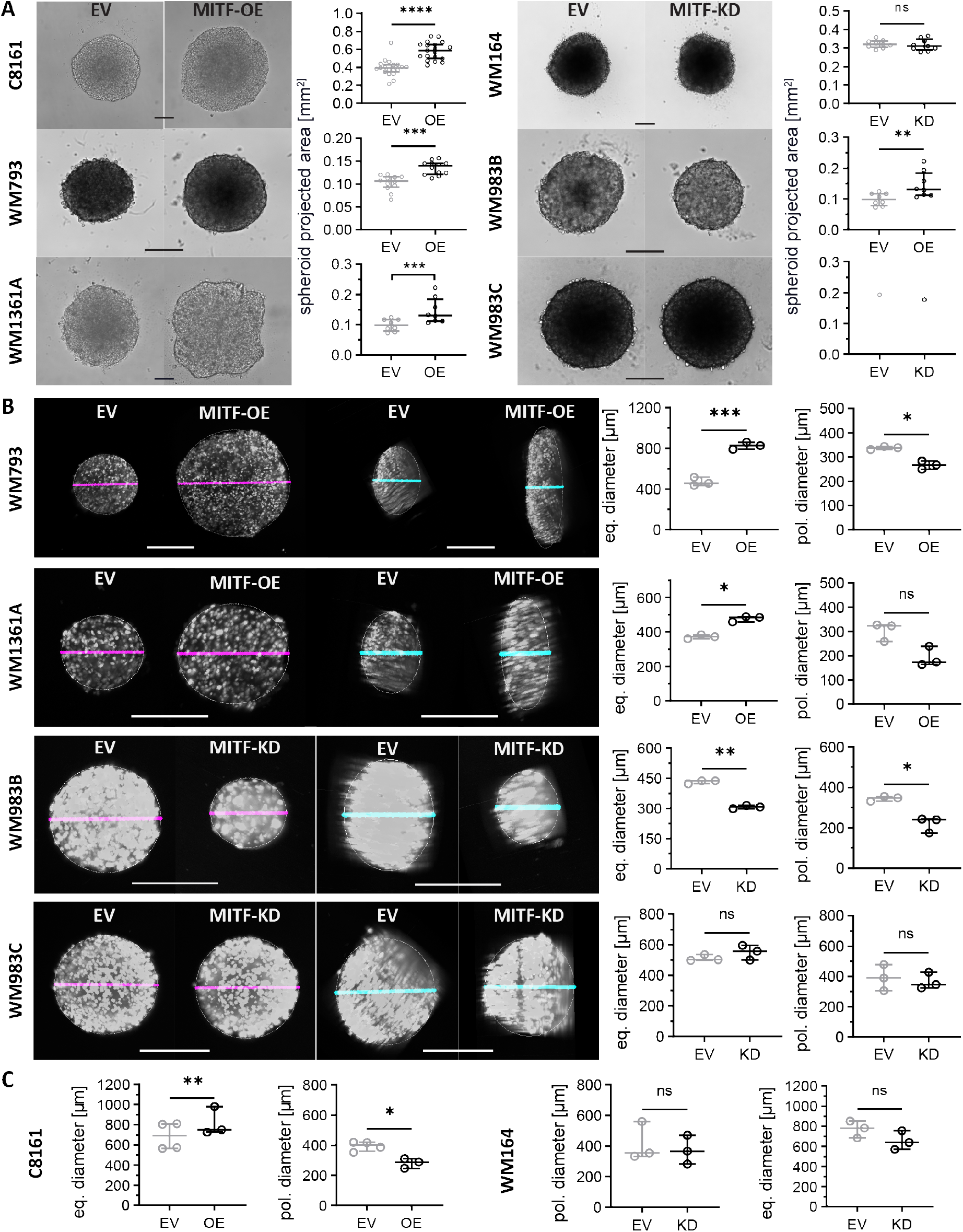
Spheroid morphology and cell count. **(A)** Representative bright-field microscopy images of spheroids and quantitation of their projected areas. **(B)** Representative SPIM images (utilizing mAG for fluorescence) and quantitation of polar and equatorial diameters of live WM793B^EV/MITF-OE^, WM1361A^EV/MITF-OE^, WM983B^EV/MITF-KD^ and WM983C^EV/MITF-KD^ spheroids. Magenta and cyan lines indicate measured equatorial and polar diameters, respectively, scale bars: 200 μm. Scatter dot plots of median and interquartile range; analysis: paired t-test, ****p<0.0001, ***p<0.001, ** p<0.01, * p<0.05, n.s. p>0.05.

**Figure S 4:**
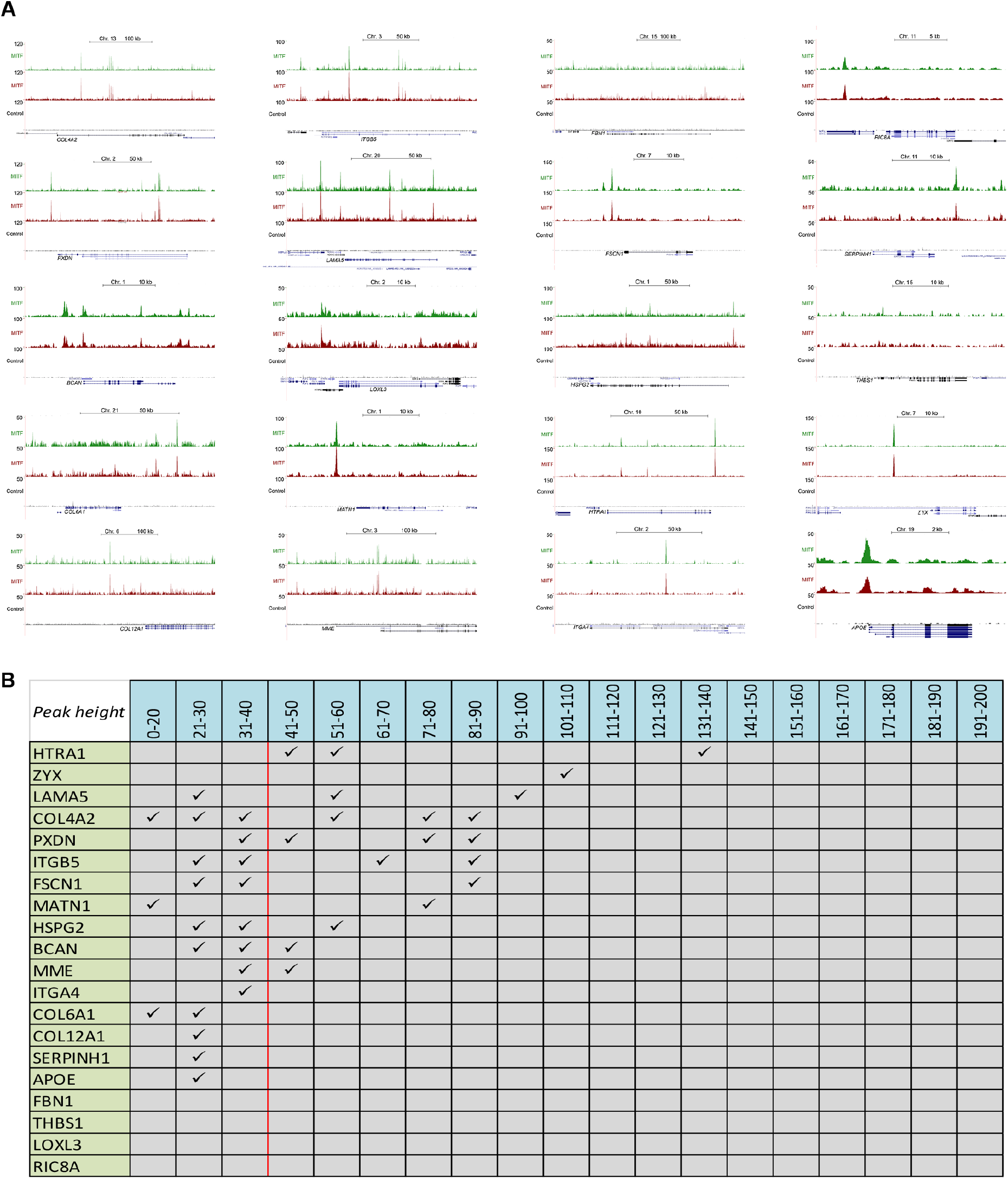
MITF ChIP-seq analysis. **(A)** UCSC browswer screen shots at indicated gene loci obtained from a published MITF ChIP-seq dataset from Louphrasitthiphol et al 2020 deposited in the Gene Expression Omnibus under accession number GSE77437. Red and green profiles indicate data from duplicate ChIP-seq experiments. **(B)** Table summarizing the ChIP-seq data shown in (A). The vertical red line indicates the cut-off for binding.

**Figure S 5:**
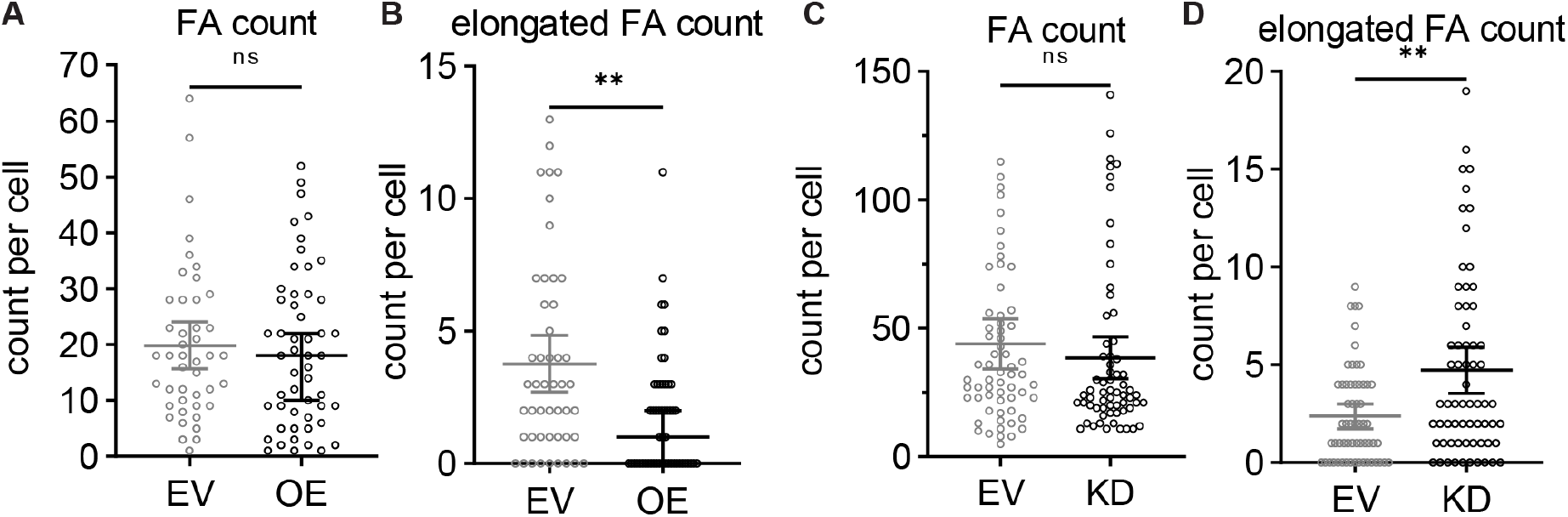
Count of focal adhesions. C8161^EV^, C8161^MITF-OE^, WM164^EV^ and WM164^MITF-KD^ cells were stained with an antibody against pFAK-Y397 and focal adhesions counted using CellProfiler software **(A and C)** and elongated focal adhesions counted manually **(B and D)**. Graph shows number of adhesions per cells (each circle represents a cell). Scatter dot plot of data median with 95% CI; each dot represents a cell; n>44 cells from three independent experiments; analysis: Mann Whitney test, ***p<0.001, ** p<0.01, n.s. p ≥ 0.05.

**Figure S 6:**
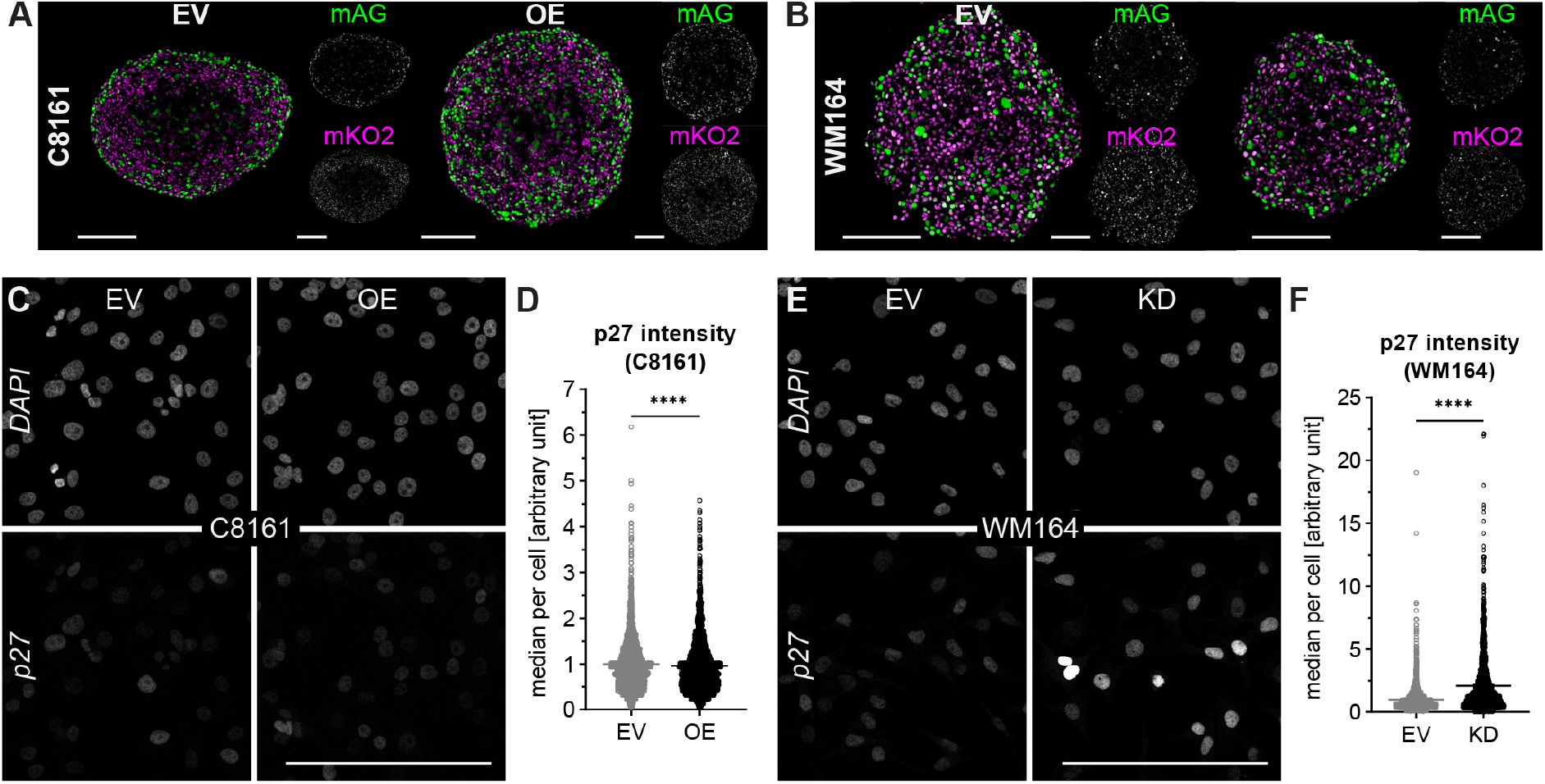
p27Kip1 protein level. **(A,B)** Corresponding FUCCI images of DAPI and p27^kip^ images shown in Figure 6A,C. **(C,E)** Representative immuno-fluorescence microscopy images of C8161 (C) and WM164 (E) adherent cultures stained with anti-p27^Kip1^ antibody. **(D,F)** Scatter dot plot of median intensity of p27^Kip1^ fluorescent signal for each cell (dots) from three independent repeats of the same experiment. Bars represent data mean with 95% CI; analysis: Mann Whitney test, **** p<0.0001. Scale bars: 200 μm.

**Figure S 7:**
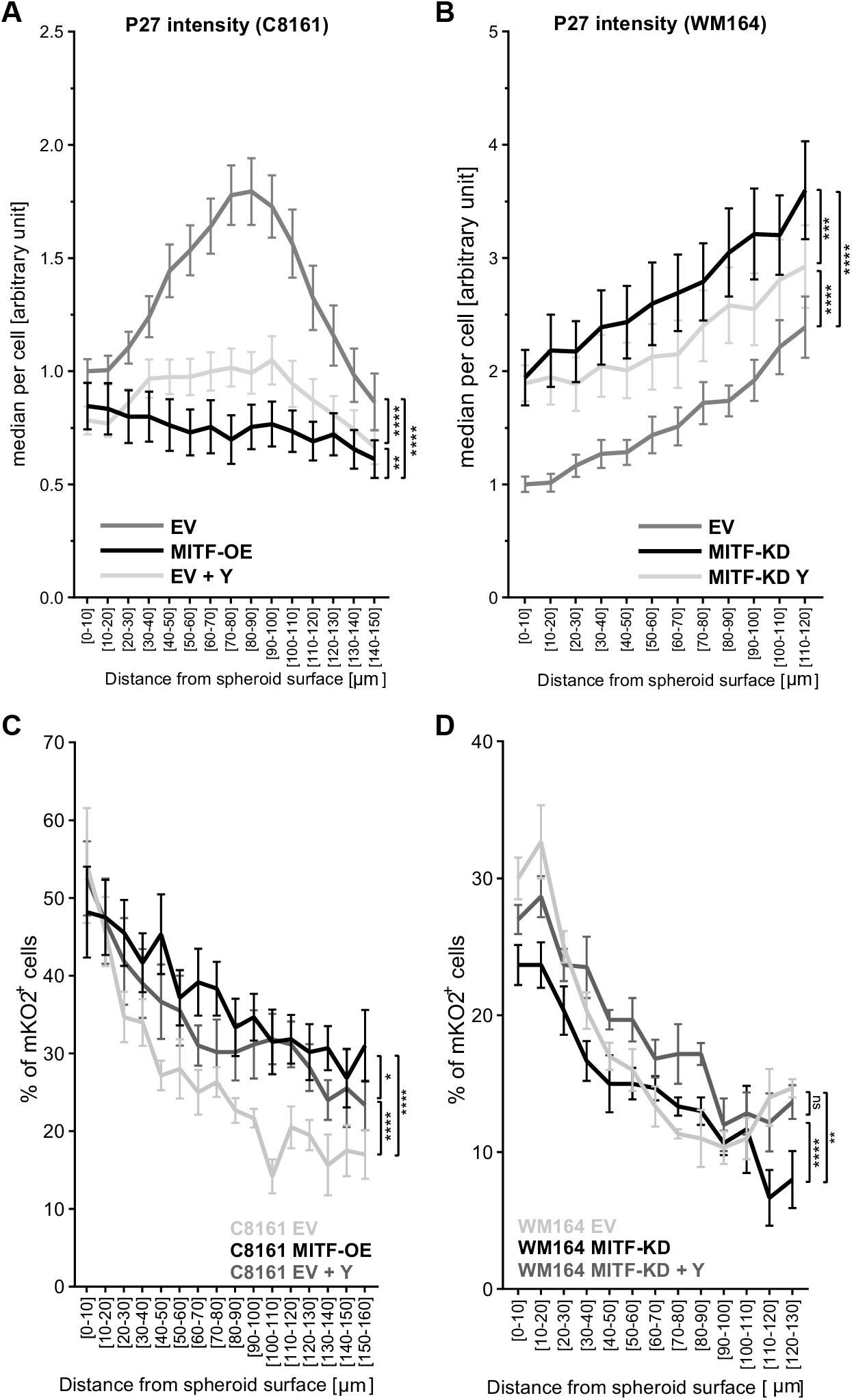
FUCCI and p27^Kip1^ immunofluorescence quantitation and analysis of ROCKi-treated spheroids. **(A,B)** Cell median p27^Kip1^ intensity plotted as a function of radial distance from the spheroid surface for the experiment described in Figure 7E. Data: mean ±SEM with two-way ANOVA analysis for the effect of MITF or Y27632 on cell median p27^Kip1^ intensity. C8161: n=21 (EV), n=15 (MITF-OE), n=13 (EV+Y); WM164: n=9 (EV), n=11 (MITF-KD), n=10 ((MITF-KD+Y); **** p<0.0001, *** p<0.001, p<0.01**. **(C,D)** Percentage of mAG^+^ cells (mAG^+^ cells/(mAG^+^ cells+ mKO2^+^ cells)) plotted as a function of radial distance from the spheroid surface for the experiment described in Figure 7F. Data: mean ±SEM with two-way ANOVA analysis for the effect of MITF or Y27632 on % of mAG+ cells. N ≥ 3; **** p<0.0001, *** p<0.001**, p<0.01, * p<0.05, ns, p≥0.05.

## Supplementary Resources

| REAGENT or RESOURCE | SOURCE | IDENTIFIER |
| --- | --- | --- |
| **Cell line** |  |  |
| 1205Lu-C5 (FUCCI) | Haass et al, 2014 | https://doi.org/10.1111/pcmr.12274 |
| 451Lu-Z1 (FUCCI) | Haass et al, 2014 | https://doi.org/10.1111/pcmr.12274 |
| C8161-A7 (FUCCI) | Haass et al, 2014 | https://doi.org/10.1111/pcmr.12274 |
| WM164-F11 (FUCCI) | Haass et al, 2014 | https://doi.org/10.1111/pcmr.12274 |
| WM793 G10 (FUCCI) | Haass et al, 2014 | https://doi.org/10.1111/pcmr.12274 |
| WM983B-X2 (FUCCI) | Haass et al, 2014 | https://doi.org/10.1111/pcmr.12274 |
| WM983C-Y3 (FUCCI) | Haass et al, 2014 | https://doi.org/10.1111/pcmr.12274 |
| **Experimental Models: Cell lines** |  |  |
| C8161<sup>EV</sup> | This paper | N/A |
| C8161<sup>MITF-OE</sup> | This paper | N/A |
| WM793<sup>EV</sup> | This paper | N/A |
| WM793<sup>MITF-OE</sup> | This paper | N/A |
| WM164<sup>EV</sup> | This paper | N/A |
| WM164<sup>MITF-KD</sup> | This paper | N/A |
| WM983B<sup>EV</sup> | This paper | N/A |
| WM983B<sup>MITF-KD</sup> | This paper | N/A |
| **Recombinant DNA** |  |  |
| MISSION® pLKO.1-puro Empty Vector Control Plasmid DNA | Sigma-Aldrich | Cat# SHC001 |
| MISSION® pLKO.1-puro Non-Mammalian shRNA Control Plasmid DNA | Sigma-Aldrich | Cat# SHC002 |
| pLVX-MITF-M plasmid DNA | This paper | N/A |
| **Viral constructs and vectors** |  |  |
| MITF MISSION® shRNA Lentiviral Transduction Particles; TRCN0000019119 (Sh19) | Sigma-Aldrich | Cat# SHCLNV-NM_000248 |
| MITF MISSION® shRNA Lentiviral Transduction Particles; TRCN0000019122 (Sh22) | Sigma-Aldrich | Cat# SHCLNV-NM_000248 |
| MITF MISSION® shRNA Lentiviral Transduction Particles; TRCN0000019123 (Sh23) | Sigma-Aldrich | Cat# SHCLNV-NM_000248 |
| Lenti-X™ HTX Packaging Mix | TaKaRa (Clontech) | Cat# 631248 |
| **Reagents** |  |  |
| MCDB 153 Medium powder | Sigma-Aldrich | Cat# M7403 |
| L-15 medium | Sigma-Aldrich | Cat# L1518 |
| Calcium Chloride | Sigma-Aldrich | Cat# 5670 |
| Insulin | Sigma-Aldrich | Cat# I0516 |
| Sodium Bicarbonate | Thermo Fisher scientific | Cat# 25080-094 |
| Sodium Hydroxide | Chem-Supply | Cat# SA178 |
| Fetal Bovine Serum | Thermo Fisher scientific | Cat# 10099141 |
| Puromycin | Sigma-Aldrich | Cat# P8833 |
| Collagen | In vitro technologies | Cat# 3442-050-01 |
| 10XMEM | Lonza | Cat# BE12-684F |
| GlutaMAX™ | Thermo Fisher scientific | Cat# 35050061 |
| Agarose | Lonza | Cat# 50004 |
| CellTracker™ Deep Red Dye | Thermo Fisher scientific | Cat# C34565 |
| Y-27632 | In vitro technologies | Cat# RDS125410 |
| Paraformaldehyde | Pro Sci Tech | Cat# C004 |
| Tissue Freezing Medium | Decision One | Cat# TFM-5-Green |
| Protease inhibitor cocktail | Sigma-Aldrich | Cat# 11836170001 |
| Phenylmethylsulfonyl fluoride | Sigma-Aldrich | Cat# 10837091001 |
| Sodium orthovanadate | Sigma-Aldrich | Cat# S6508-10G |
| Sodium fluoride | Sigma-Aldrich | Cat# S7920-100G |
| Aprotinin | Sigma-Aldrich | Cat# 10236624001 |
| Leupeptin | Sigma-Aldrich | Cat# L2884-1MG |
| NuPAGE™ 4-12% Bis-Tris Protein Gels | Thermo Fisher scientific | Cat# NP0321BOX |
| Immobilon-P PVDF Membrane | Millipore | Cat# IPVH00010 |
| Immun-Blot® Low Fluorescence PVDF membrane | BIO-RAD | Cat# 1620264 |
| Clarity™ Western ECL Substrate | BIO-RAD | Cat# 1705060 |
| DTT | Astral Scientific | Cat# C-1029-25g |
| C18 tips | Glygen | Cat# Velo-C18-20ug |
| Column Easy-Spray 500 x 0.075mm ID 2um | Thermo Scientific | Cat# THCES803A |
| **KITS** |  |  |
| Pierce™ BCA Protein Assay Kit | Thermo Fisher scientific | Cat# 23227 |
| **Software and Algorithms** |  |  |
| Cell profiler | Broad Institute | cellprofiler.org/ |
| ImageJ | National Institute of Health | imagej.nih.gov/ij/ |
| GraphPad Prism | GraphPad Software | www.graphpad.com/scientific-software/prism/ |
| Volocity | PerkinElmer | www.perkinelmer.com/volocity |
| R version 3.6 | The R Project for Statistical Computing | https://www.r-project.org/ |
| Pheatmap | N/A | https://CRAN.R-project.org/package=pheatmap |
| PicardTools version 1.96 | N/A | http://picard.sourceforge.net |
| Bowtie 1.1.2 | (102) | N/A |
| **Antibodies** |  |  |
| Anti-MITF | Agilent Technologies | Cat# M362129-2; RRID:AB_2142100 |
| Anti-Rab27a | Sigma-Aldrich | Cat# WH0005873M2; RRID:AB_1843221 |
| Anti-GAPDH | R and D Systems | Cat# 2275-PC-020; RRID:AB_2107456 |
| Anti-pMLC | Cell Signaling Technology | Cat# 3671S; RRID:AB_330248 |
| Anti-pFAK | Cell Signaling Technology | Cat# 8556S; RRID:AB_10891442 |
| Anti-Mouse IgG-HRP | Invitrogen | Cat# 626520; RRID:AB_88369 |
| Anti-Rabbit IgG-HRP | Cell Signaling Technology | Cat# 7074S; RRID:AB_2099233 |
| Anti-Mouse IgG- Alexa Fluor 647 | Thermo Fisher Scientific | Cat# A21236; RRID:AB_141725 |
| Anti-Rabbit IgG- Alexa Fluor 647 | Thermo Fisher Scientific | Cat# A31573; RRID:AB_2536183 |
| Anti-Rabbit IgG- Alexa Fluor 750 | Abcam | Cat# ab175729; |

## Lead Contact and Material Availability

Request for further information and resources should be directed to Nikolas Haass

The cell lines generated in this paper can be made available with approved material transfer agreements from The University of Queensland, The Centenary Institute, The Wistar Institute and RIKEN.

